# Disentangling the flow of signals between populations of neurons

**DOI:** 10.1101/2021.08.30.458230

**Authors:** Evren Gokcen, Anna I. Jasper, João D. Semedo, Amin Zandvakili, Adam Kohn, Christian K. Machens, Byron M. Yu

## Abstract

Technological advances now allow us to record from large populations of neurons across multiple brain areas. These recordings may illuminate how communication between areas contributes to brain function, yet a substantial barrier remains: How do we disentangle the concurrent, bidirectional flow of signals between populations of neurons? We therefore propose here a novel dimensionality reduction framework: Delayed Latents Across Groups (DLAG). DLAG disentangles signals relayed in each direction, identifies how these signals are represented by each population, and characterizes how they evolve within and across trials. We demonstrate that DLAG performs well on synthetic datasets similar in scale to current neurophysiological recordings. Then we study simultaneously recorded populations in primate visual areas V1 and V2, where DLAG reveals signatures of bidirectional yet selective communication. Our framework lays a foundation for dissecting the intricate flow of signals across populations of neurons, and how this signaling contributes to cortical computation.

## Introduction

Simultaneous recordings from large populations of neurons across multiple brain areas are growing in availability [1–4]. These recordings present opportunities to illuminate how inter-areal communication enables brain function [5], but they also present substantial conceptual and statistical challenges. Brain areas involved in sensory [6–9], cognitive [10], and motor functions [11] are often reciprocally connected: signals are relayed not only from one area to the next, but bidirectionally, and likely concurrently. The raw recordings, however, provide only a tangled view of this concurrent communication (Fig. 1, top): individual neurons simultaneously reflect an area’s inputs, outputs, and ongoing internal computations [12].

**Figure 1.**
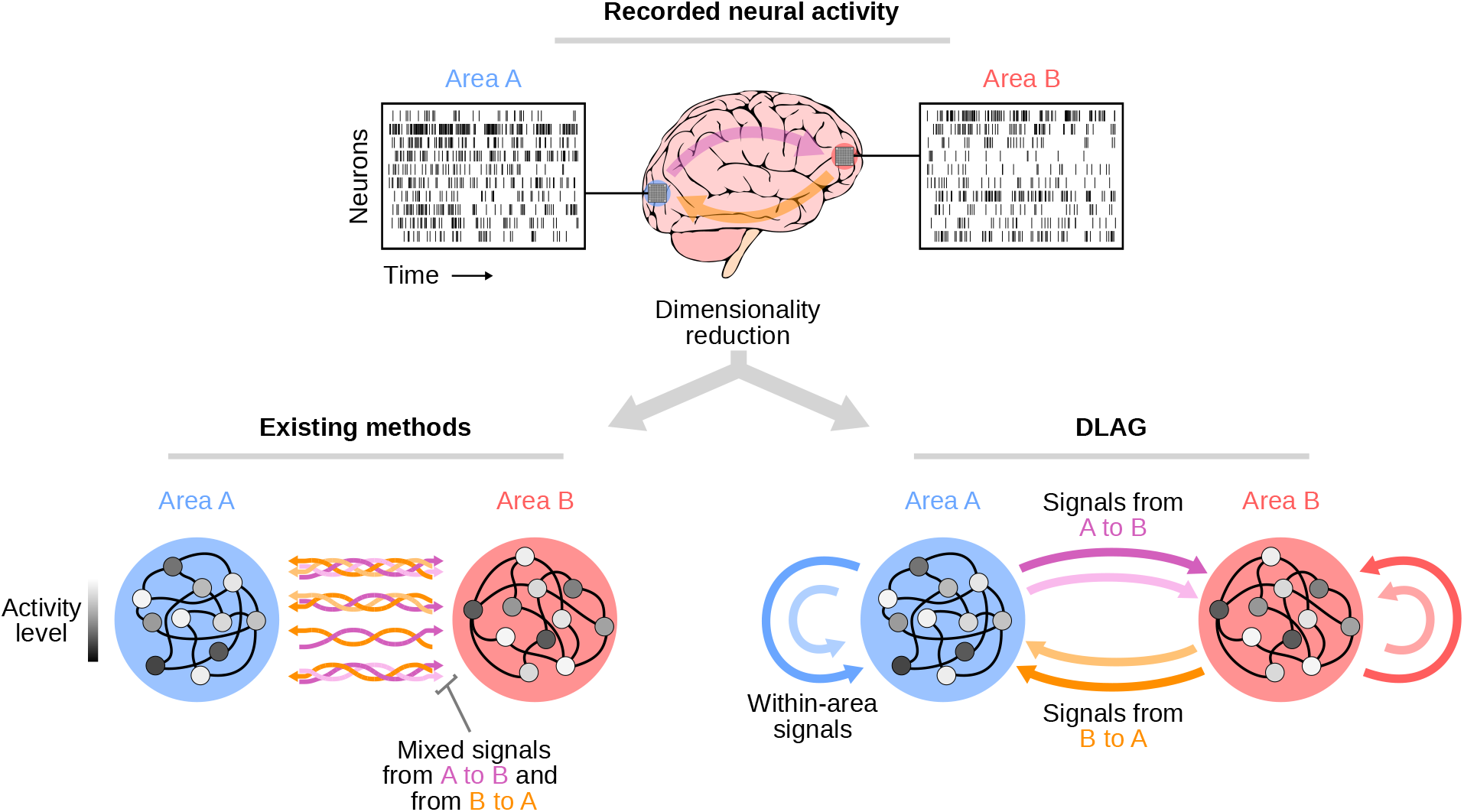
Disentangling the flow of signals between populations of neurons. Top: Recorded neural activity provides only a tangled view of the bidirectional, concurrent interactions between brain areas (illustrated by the thick translucent arrows; magenta: signals directed from area A to area B; orange: signals directed from area B to area A). Bottom left: Existing dimensionality reduction methods identify correlated population activity across areas (each correlated population activity pattern is represented by a braid of multi-colored arrows; four different activity patterns are shown). Each activity pattern likely reflects a mixture of signals relayed in each direction. Within each activity pattern, individual arrows represent a directed interaction; color depicts the direction of signal flow (magenta: A to B; orange: B to A), and shading (light vs. dark) distinguishes distinct signals. Bottom right: We propose a novel dimensionality reduction method, Delayed Latents Across Groups (DLAG). DLAG identifies both within-and across-area population signals (indicated by color and source/target of each arrow; blue: within-A; red: within-B; magenta/orange: across-area). Importantly, DLAG disentangles signals relayed in each direction. The color of each arrow depicts the direction of signal flow (magenta: A to B; orange: B to A) associated with a population activity pattern, and shading (light vs. dark) distinguishes distinct signals.

Determining the flow of signals between brain areas is therefore a nontrivial task. To dissect the direction of signal flow, one can leverage the fact that inter-areal communication is not instantaneous. The physiological properties of axons and synapses introduce delays in signal transmission. These delays provide a working definition of signal flow: the appearance of a signal first in area A, and later in area B, is consistent with signal flow from A to B (though this apparent flow could be due to common input from a third area; see Discussion).

Adopting this conception, several inter-areal studies have compared the timing of the onset of neural responses [13–16] or of the emergence of selectivity attributable to top-down processes [17–21] across areas following the presentation of a stimulus. Other studies, leveraging simultaneous recordings, have measured temporal delays between two areas through pairwise spiking correlations [22–27] and information theoretic measures [28, 29]. Similarly, inter-areal phase delays of local field potentials (LFPs) have been measured [30–33]. These timing-based approaches have significantly advanced our understanding of how signals propagate across brain areas. However, given that neuronal computations are believed to be carried out by neuronal populations, such approaches—focused largely on pairs of neurons or aggregate measures of neural activity—lack the richness to fully describe the nature of signals relayed between areas.

A full characterization of inter-areal signal flow therefore requires relating the activity of populations of neurons across two or more areas—a challenging high-dimensional problem. Dimensionality reduction techniques capable of identifying low-dimensional latent variables that describe activity shared by two or more recorded areas are thus increasingly used [34–36]. These techniques have driven new proposals for population-level mechanisms of gating between motor cortex output and muscle movement [37, 38]; selective communication between cortical areas [39, 40]; enhanced communication of stimulus information with attention [41]; and the robustness of local computations to perturbations upstream [42, 43].

The relationship between the correlated activity across areas identified in these studies and the flow of inter-areal signals, however, remains unclear. Specifically, does the correlated activity across areas reflect the flow of activity from area A to B, from B to A, or in both directions concurrently? If communication were to occur in one direction at a time, then existing dimensionality reduction methods could, in principle, identify the direction of population-level signal flow. If two areas were to communicate in both directions concurrently, however, then existing methods would only identify the dominant direction of signal flow [44]. Disentangling the concurrent flow of signals between populations remains a substantial barrier in neuroscience (Fig. 1, bottom left).

We therefore propose a novel dimensionality reduction framework: Delayed Latents Across Groups, or DLAG (Fig. 1, bottom right). DLAG disentangles signals relayed in each direction, identifies how these signals are represented by each population, and characterizes how they evolve within and across trials. We first demonstrate that DLAG performs well on synthetic datasets similar in scale to current neurophysiological data. Then we study simultaneously recorded populations in primate visual areas V1 and V2, where DLAG reveals that V1-V2 interactions are selective and bidirectional. DLAG unlocks new opportunities to investigate the bidirectional flow of signals between populations of neurons and how inter-areal communication contributes to brain function.

## Results

### Delayed Latents Across Groups (DLAG)

Consider recording the activity of two populations of neurons (Fig. 2, left column), measured as, for example, the number of spikes counted within nonoverlapping time bins. Here we will take these populations as belonging to two different brain areas, A and B. In principle, they can belong to any meaningful groups, such as cortical layers or cell types.

**Figure 2.**
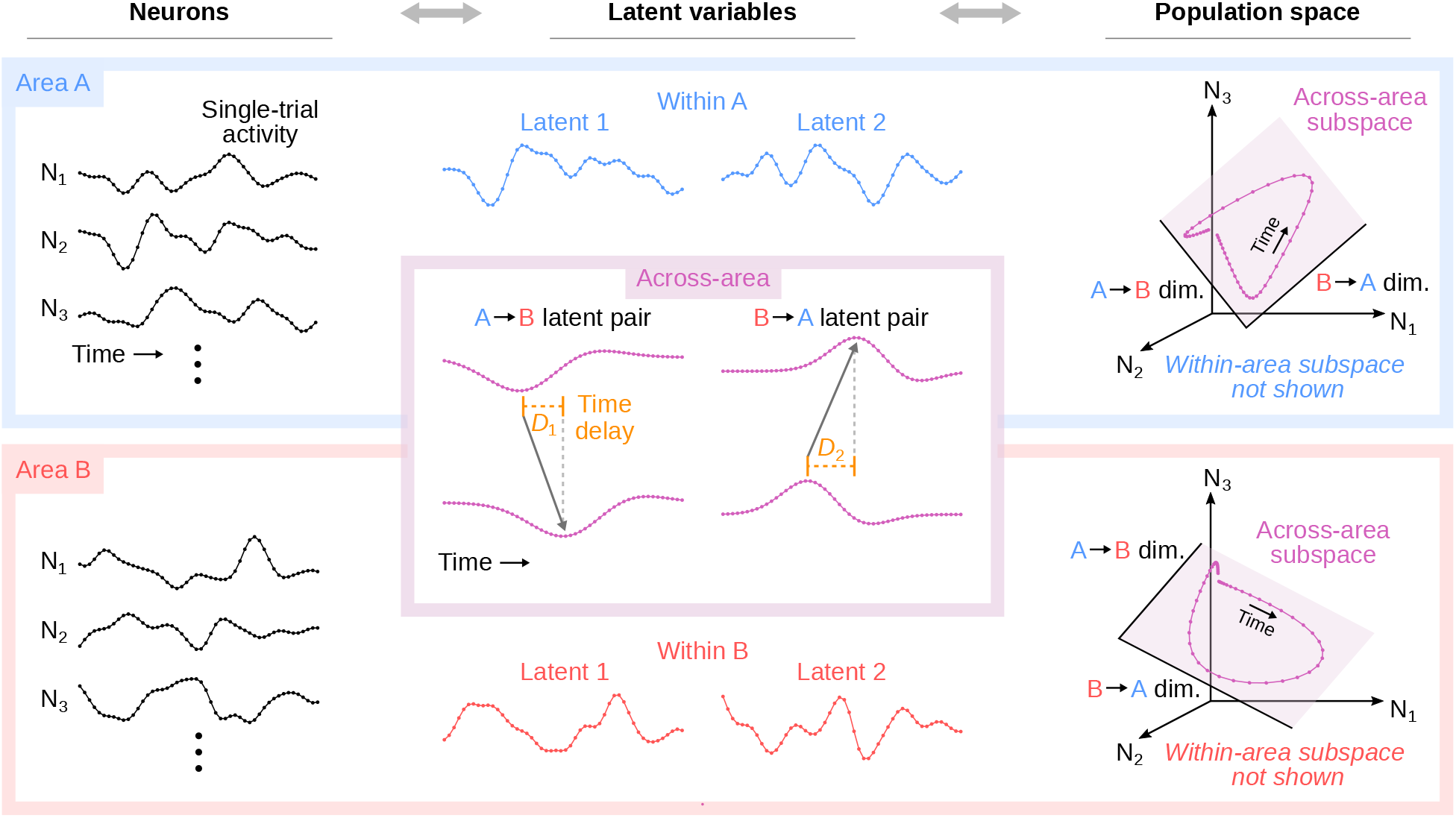
DLAG conceptual illustration. From left to right: neurons, latent variables, and population activity space representations in two recorded brain areas analyzed by DLAG (top row / blue box: area A; bottom row / red box: area B). Left column: Single-trial activity of neurons simultaneously recorded in each area. Only three neurons (N_1_, N_2_, N_3_) are shown in each area for clarity. Center column: DLAG expresses the observed neural activity in each area as a linear combination of two types of low-dimensional latent variables: within-area and across-area. Within-area variables are shown in the color corresponding to the area in which they belong (Within A: blue; Within B: red). For clarity, only two within-area variables are shown in each area, but in principle there may be a greater number, as determined by DLAG from the recorded activity. Across-area variables are shown in magenta. The magenta box inset overlaps the blue and red boxes for area A and B, respectively, to indicate that across-area variables are shared among neurons in both areas. The location of each across-area variable (i.e., within the bounds of Area A’s box or area B’s box) indicates which area’s activity it reflects. Between area A and area B, across-area variables are vertically paired. The time courses of each pair are related after a time delay (*D*_1_: delay between the left pair; *D*_2_: delay between the right pair). The sign of this delay allows each pair to be associated with a directed interaction (A to B or B to A), which is indicated by gray arrows. For clarity, only two across-area variable pairs are shown. Right column: The activity of each neural population can be represented in a population activity space, where each axis represents the activity of a single neuron (N_1_, N_2_, N_3_). Each point in population space represents the population activity at a particular time, and the points trace out a trajectory over time (magenta curve). DLAG identifies two linearly independent subspaces in each area: a within-area subspace (not shown, for clarity) and an across-area subspace (magenta-shaded plane). Each dimension (‘dim.’) of the across-area subspace is associated with a directed interaction.

DLAG dissects the recorded population activity in each area on individual trials into a linear combination (weighted sum) of two types of latent variables (Fig. 2, center column). The first type of latent variable, *across-area* variables, describes population activity that is correlated across areas (illustrated by the magenta box spanning both areas in Fig. 2). The second type of latent variable, *within-area* variables, describes population activity in one area that is not related to population activity in the other area (Fig. 2; blue: within A; red: within B). Whether or not the within-area variables are a subject of scientific study, they are critical to the correct estimation of across-area variables (see Methods and Supplementary Discussion).

Across-area variables are defined in pairs, where the elements of each pair correspond to the two areas. Importantly, the elements of each pair are time-delayed relative to each other (Fig. 2, *D*_1_ between the first pair and *D*_2_ between the second pair). Consequently, if a particular time course is reflected in the population activity of area A, and a similar time course is reflected in the population activity of area B, but after a time delay, then an across-area variable pair can describe the apparent flow of that signal from A to B. And if, concurrently, a time course is first seen in area B, followed by area A, a second across-area variable pair can also describe the flow of that inter-areal signal. The key to disambiguating the first and second across-area variable pairs is that they involve different population activity patterns (i.e., a “loading” vector indicating how the activity of each neuron relates to the latent variable). In fact, DLAG can identify many across-area variable pairs, each with a delay of its own sign and magnitude, to capture multiple concurrent streams of signal flow between the two populations at different timescales.

The relationship between within- and across-area latent variables and observed population activity in each area can be represented geometrically with the concept of a population activity space (Fig. 2, right column). For each area, we can define a high-dimensional population activity space where each axis represents the activity of one neuron. Each point in the space represents the population activity at a particular time, and the points trace out a trajectory over time. DLAG’s two types of latent variables each define the axes (dimensions) of a low-dimensional sub-space within this population activity space (in Fig. 2, we show only the across-area subspaces for visual clarity). Each dimension of these subspaces represents a population activity pattern.

The temporal structure of within- and across-area variables are both described by relating each latent variable at different time points through Gaussian processes (see Methods and Supplementary Fig. 1). Each Gaussian process is associated with its own characteristic timescale that controls the temporal smoothing of neural activity (Supplementary Fig. 2). DLAG estimates both these timescales and time delays from the neural activity using an exact expectation-maximization (EM) algorithm. After the DLAG model parameters are estimated from the neural activity, the time courses of within- and across-area latent variables can be studied on a trial-to-trial basis.

### Validation on realistic-scale synthetic data

Before applying DLAG to experimental data, we characterized its performance on synthetic datasets similar in scale to state-of-the-art neurophysiological recordings from multiple brain areas, and on additional synthetic datasets covering a wider range of experimental conditions. Informed by our recordings in macaque V1 and V2 [27, 39] (see next section), we simulated independent datasets with representative numbers of neurons (area A: 80; area B: 20), trial counts (100), trial lengths (1,000 ms), and levels of noise, where noise is defined as the variance independent to each neuron (see Methods for additional details).

#### Estimation of parameters and latent variables

Across all datasets, within- and across-area latent time courses (Fig. 3a; see legend for quantification), across-area parameters (Fig. 3b, dimensionalities; Fig. 3c, delays; Fig. 3d, Gaussian process timescales), and within-area parameters (Fig. 3e, dimensionalities; Fig. 3f,g, Gaussian process timescales) were all consistently and accurately estimated. We highlight, in particular, DLAG’s ability to estimate time delays between the two areas (Fig. 3c). Delay error was 1.3±0.1 ms (mean and SEM across all delays; max error 7.0 ms), despite observations occurring at 20 ms time steps. This accuracy emphasizes an important feature of the DLAG model that distinguishes it from other time series modeling approaches (see Discussion). Because latent time courses and time delays are continuous-valued, DLAG can leverage the correlated activity of the neuronal populations to recover delays that are smaller than the sampling period (i.e., spike count bin width, in the case of spiking activity).

**Figure 3.**
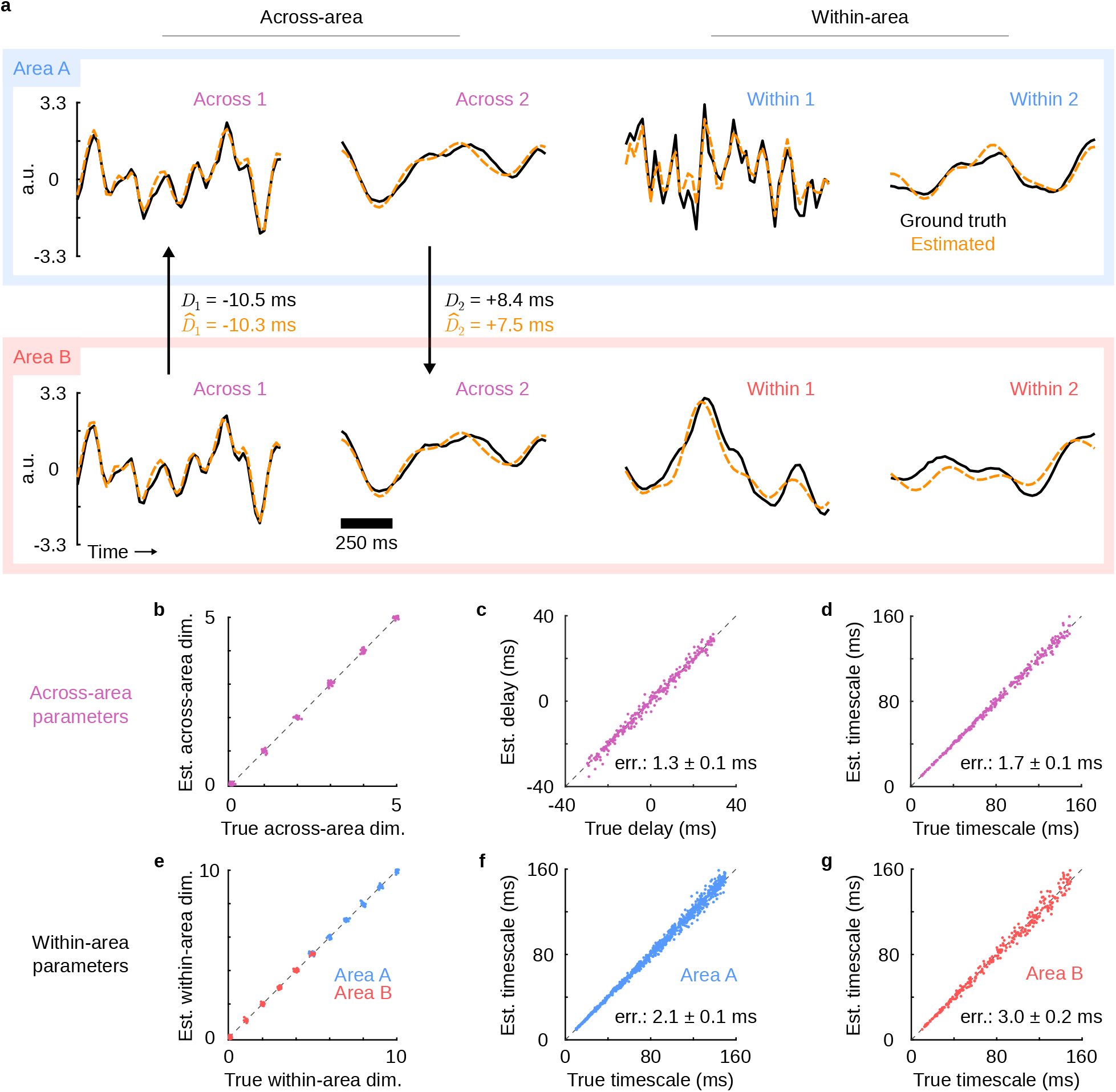
DLAG accurately estimates within- and across-area time courses and their parameters in synthetic data. (**a**) Single-trial latent-variable time course estimates for a representative synthetic dataset. Top row / blue box: area A; bottom row / red box: area B. For visual clarity, two latent variables of each type are shown (left: across-area; right: within-area). Orange dashed traces: DLAG estimates; black solid traces: ground truth. a.u.: arbitrary units. Across all synthetic datasets for which across- or within-area dimensionality was non-zero (see Methods; across: 100 datasets; within A: 120 datasets; within B: 100 datasets), mean accuracy (*R*^2^) of time course estimation was as follows: area A, across − 0.90; area B, across − 0.91; area A, within − 0.88; area B, within − 0.82 (all SEM values less than 0.01). Similarly, mean accuracy of subspace (loading matrix) estimation was as follows: 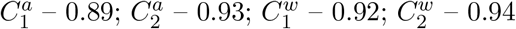 (where a value of 1 implies that the ground truth is fully captured by estimates; all SEM values less than 0.01). (**b**) Across-area dimensionality estimates versus the ground truth for all 120 synthetic datasets. Data points are integer-valued, but randomly jittered to show points that overlap. (**c**) Delay estimates versus the ground truth. Displayed error (‘err.’) indicates mean absolute error and SEM reported across 300 across-area variables. (**d**) Across-area Gaussian process (GP) timescale estimates versus the ground truth. Displayed error (‘err.’) indicates mean absolute error and SEM reported across 300 across-area variables. (**e**) Within-area dimensionality estimates versus the ground truth for all 120 synthetic datasets (blue: within-area A; red: within-area B). Data points are integer-valued, but randomly jittered to show points that overlap. (**f**) Within-area A GP timescale estimates versus the ground truth. Displayed error (‘err.’) indicates mean absolute error and SEM reported across 900 within-area variables in area A. (**g**) Within-area B GP timescale estimates versus the ground truth. Displayed error (‘err.’) indicates mean absolute error and SEM reported across 300 within-area variables in area B.

The synthetic datasets presented here were generated with a variety of parameters representative of realistic data, but we also verified that DLAG performed well over a wider range of simulated conditions. Specifically, we systematically characterized DLAG’s performance as a function of number of trials (Supplementary Fig. 3), number of neurons (Supplementary Fig. 4), and latent timescale (Supplementary Fig. 5). We also characterized the runtime of the DLAG fitting procedure as a function of number of trials, number of neurons, trial length, and latent dimensionality (Supplementary Fig. 6).

#### Scalable model selection

Estimating the number of within- and across-area latent variables is a challenging problem. For example, performing a grid search over just 10 possibilities for each type of latent variable (within-area A, within-area B, and across-area) would result in 1,000 model candidates. We therefore developed a streamlined cross-validation procedure that significantly improves scalability (see Methods). In brief, we apply factor analysis [45] to each area separately to estimate the dimensionality of population activity in that area. We then use these estimates to constrain the total number of within- and across-area latent variables.

Applied to the synthetic datasets described above, our cross-validation procedure proved highly accurate. Across all datasets—including those with no across- or within-area structure—the selected dimensionalities matched the ground truth (Fig. 3b, across-area; Fig. 3e, within-area). Furthermore, model selection remained accurate in additional, more challenging scenarios, where we considered synthetic datasets with significantly lower signal-to-noise ratios (lower than typically encountered in our V1-V2 recordings, below) (Supplementary Fig. 7). And, in the instances where the greater statistical challenge induced imperfect estimates of dimensionality, DLAG’s parameter and latent variable estimates remained stable (Supplementary Fig. 8, Supplementary Fig. 9).

### Dissecting bidirectional interactions between V1 and V2

We then used DLAG to study interactions between two areas in the early visual system: V1 and V2. V1 and V2 share strong reciprocal connections [46, 47] and show correlated activity [23–25, 27, 39], but the bidirectional nature of their interactions is not yet well understood. We simultaneously recorded the activity of neuronal populations in the superficial (output) layers of V1 (61 to 122 neurons; mean 86.3), and the middle (input) layers of V2 (15 to 32 neurons; mean 19.6) in three anesthetized monkeys (Fig. 4a; data reported previously in [27, 39]). Recording locations were selected to maximize the probability that the recorded V1 and V2 populations interact by ensuring spatial receptive field alignment. We analyzed neuronal responses measured during the 1.28 second presentation of drifting sinusoidal gratings of different orientations, and counted spikes in 20 ms time bins. The periodic nature of the drifting gratings (160 ms per cycle) is evident in peristimulus time histograms (PSTHs) for an example recording session and grating orientation (Fig. 4b). In total, we fit DLAG models separately to 40 “datasets,” corresponding to five recording sessions, each with eight different orientations. For comparison, we also applied DLAG to two V1 subpopulations (termed V1a and V1b) (Fig. 4c).

**Figure 4.**
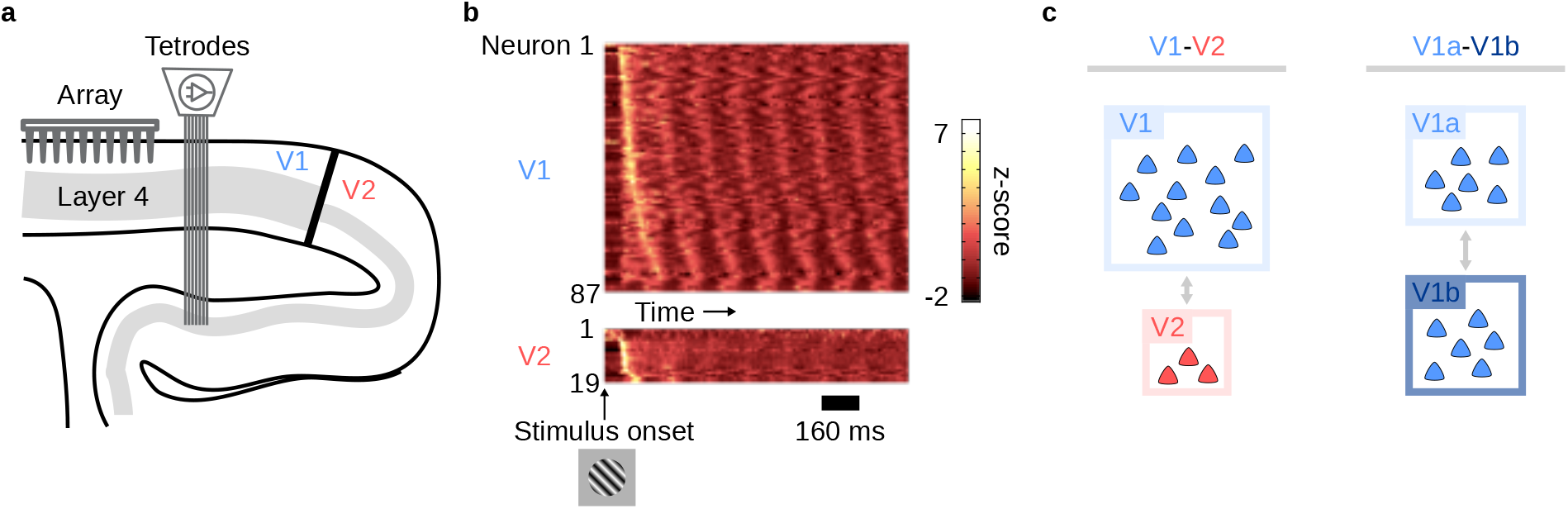
Simultaneous population recordings in V1 and V2. (**a**) Schematic showing a sagittal section of occipital cortex and the recording setup. V1 population activity was recorded using a 96-channel Utah array. V2 population activity was recorded using a set of movable electrodes and tetrodes. (**b**) Peristimulus time histograms during the stimulus presentation period, for an example session and stimulus condition. For visualization purposes, neuronal spike trains were first smoothed using a sliding Gaussian window of width 20 ms, and then z-scored to produce normalized firing rates. Neurons are ordered from top to bottom (separately for V1 and V2) according to the time at which their peak firing rate occurs. (**c**) Inter- and intra-areal comparisons. (Left) We applied DLAG to spike counts in V1 (light blue) and V2 (red). (Right) For comparison, we applied DLAG to two equally sized V1 subpopulations (V1a, light blue; V1b, dark blue), randomly selected from the V1 population. Each triangle represents a neuron. Box sizes illustrate typical relative population sizes.

#### V1-V2 interactions are selective and are more prominent in V2 than in V1

We first used DLAG to study whether V1 and V2 interact selectively: in addition to fluctuations shared between V1 and V2, are there fluctuations that are not shared between the two areas? Selective inter-areal communication may be a hallmark of cortical computation that remains to be fully understood, particularly at the level of neuronal populations [5]. Indeed, significant across- and within-area latent variables (i.e., latent variables that were selected via cross-validation) were identified consistently across datasets (Fig. 5a: single-trial latent time courses from a representative dataset; Fig. 6a, top: dimensionalities across all datasets; median dimensionality across areas: 3; within-V1: 14; within-V2: 2).

**Figure 5.**
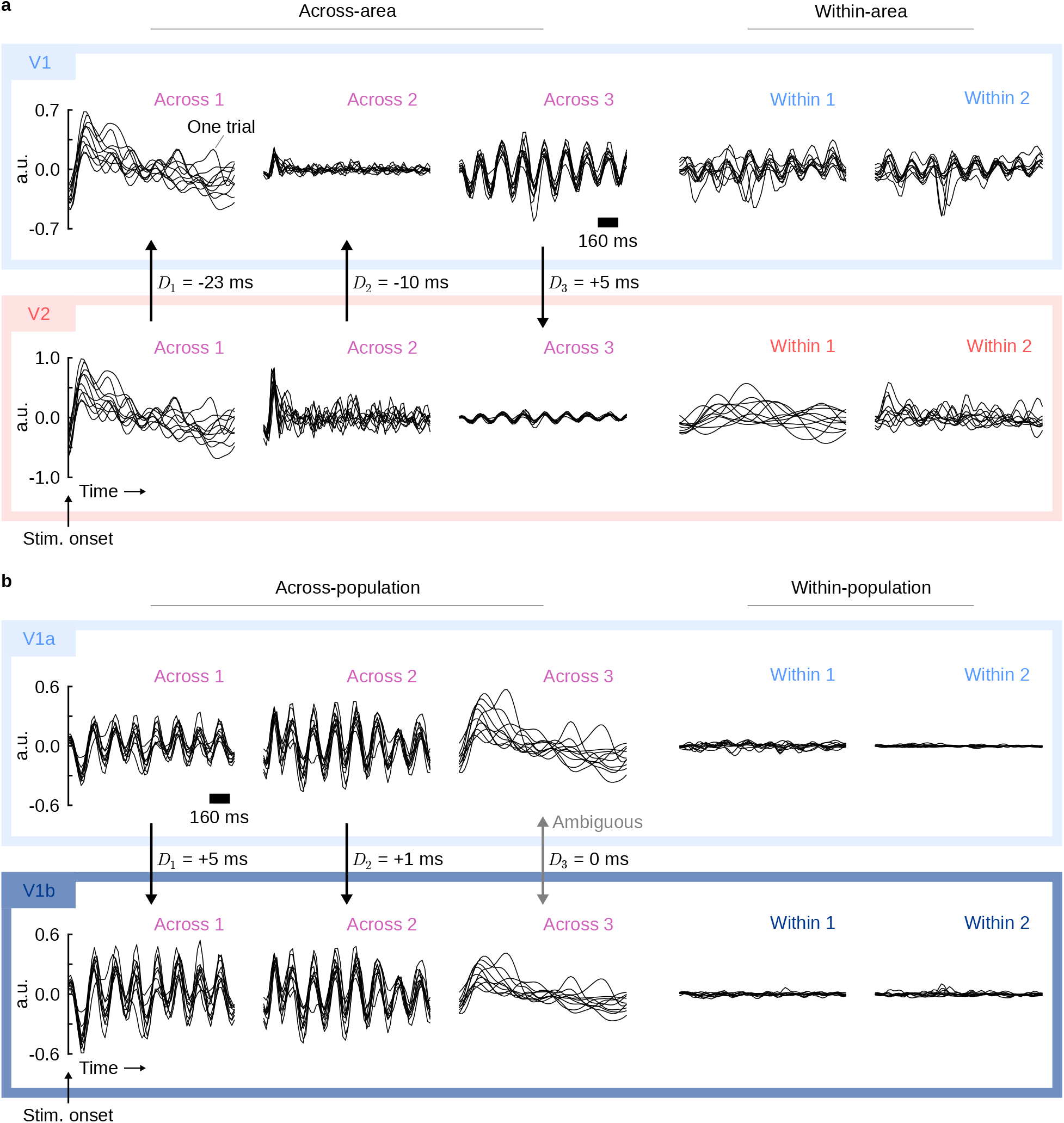
Representative DLAG time courses for inter- and intra-areal analyses (same dataset as shown in Fig. 4b). (**a**) Inter-areal (V1-V2) time courses. Left: Across-area time courses. Right: Within-area time courses. Top row / light blue box: V1. Bottom row / red box: V2. Each panel corresponds to the single-trial time courses of a latent variable. All time courses are aligned to stimulus onset. a.u.: arbitrary units. Each black trace corresponds to one trial; for clarity, only 10 of 400 are shown. Note that the polarity of traces is arbitrary, as long as it is consistent with the polarity of 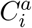 or 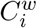. Across-area variables are paired vertically; vertical arrows point in the direction of the identified signal flow, as determined by the sign of the delay next to each arrow. All delays for the displayed dataset were deemed significantly different from zero (see Methods). For visualization purposes, latent variables have been scaled and ordered by the fraction of shared variance they explain (across- and within-area variables are sorted separately; across-area variables are sorted according to shared variance explained in V2). All across-area variables and within-V2 variables uncovered by DLAG are shown here. The top 2 of 14 within-V1 variables are displayed, which explain 46% of V1’s within-area shared variance. (**b**) Intra-areal (V1a-V1b) time courses. Left: Across-population time courses. Right: Within-population time courses. Top row / light blue box: V1a. Bottom row / dark blue box: V1b. All other conventions the same as in (a). Here, the delay for the third across-population variable (Across 3) was deemed to have an ambiguous sign, indicated by the bidirectional gray arrow. All other delays for the displayed dataset were deemed significantly different from zero, indicated by the unidirectional black arrows. Three of 10 across-population variables uncovered by DLAG are shown here, which explain 23% and 17% of V1a’s and V1b’s total shared variance, respectively. All uncovered within-V1a variables are shown, and 2 of 5 within-V1b variables are shown. Within-population variables (including those not shown here) explained 5% and 7% of V1a’s and V1b’s total shared shared variance, respectively.

**Figure 6.**
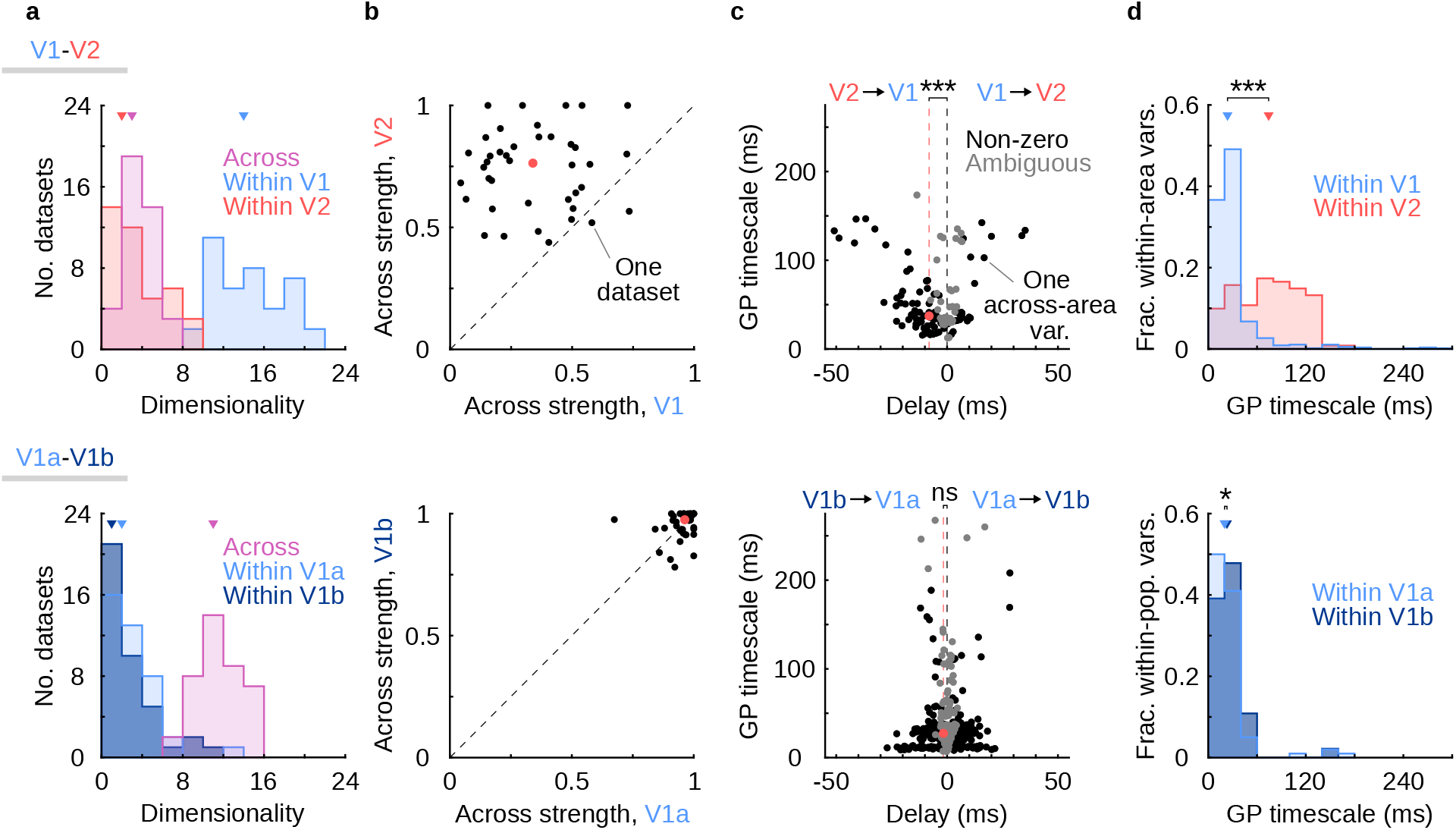
DLAG reveals that V1-V2 interactions are selective and asymmetric. Top row: Summary of V1-V2 (inter-areal) interactions. Bottom row: Summary of V1a-V1b (intra-areal) interactions. (**a**) Within- and across-area dimensionalities (determined via cross-validation). Top: V1V2 results. Distribution of within-V1 (light blue), within-V2 (red), and across-area (magenta) dimensionalities across 40 datasets. Triangles indicate the median of each distribution (within-V1: 14; within-V2: 2; across-area: 3). Within-V2 dimensionality was 0 in 5 of 40 datasets. Bottom: V1a-V1b results. Distribution of within-V1a (light blue), within-V1b (dark blue), and across-population (magenta) dimensionalities across 40 datasets. Triangles indicate the median of each distribution (within-V1a: 2; within-V1b: 1; across-population: 11). Within-V1a dimensionality was 0 in 8 of 40 datasets; within-V1b dimensionality was 0 in 11 of 40 datasets. (**b**) Fraction of shared variance of each area explained by across-area latent variables. Top: V1-V2 results. Black points: across-area strength in V1 vs. across-area strength in V2 for each of 40 datasets. Red point: median across datasets (V1: 0.34; V2: 0.76). Across-area strength is significantly greater in V2 than in V1 (one-sided paired sign test; *p* < 0.001). Bottom: V1a-V1b results. Black points: across-population strength in V1a vs. across-population strength in V1b for each of 40 datasets. Red point: median across datasets (V1a: 0.96; V1b: 0.98). Across-population strength is not significantly greater in one population or the other (two-sided paired sign test; *p* = 0.87). (**c**) Gaussian process (GP) timescale vs. time delay for across-area latent variables. Top: V1-V2 results. Each point represents one across-area latent variable. Black points: across-area latent variables for which the delays were deemed significantly non-zero (see Methods; 95 of 135 across-area variables across all 40 datasets). Gray points: across-area latent variables for which delays were deemed ambiguous (not significantly positive or negative; 40 of 135 across-area variables across all 40 datasets). Red point: median across all significantly non-zero across-area variables (GP timescale: 37 ms; Delay: −8 ms). Red dashed line indicates the median delay. ‘***’: delays are significantly less than zero, representing feedback interactions from V2 to V1 (one-sided one-sample sign test on ‘non-zero’ delays, *p* < 0.001). Bottom: V1a-V1b results. Same conventions as top panel. Out of 437 across-population latent variables uncovered across all 40 datasets, 316 delays were deemed significantly non-zero (black points), while 121 delays were deemed ambiguous (gray points). Red point: median across all significantly non-zero across-population variables (GP timescale: 27 ms; Delay: −2 ms). ‘ns’: delays are not significantly negative (one-sided one-sample sign test on ‘non-zero’ delays, *p* = 0.08). (**d**) GP timescales for within-area latent variables. Top: V1-V2 results. Normalized distribution of within-V1 (light blue) and within-V2 (red) GP timescales across all 40 datasets (total within-V1 latent variables: 562; total within-V2 latent variables: 121). Triangles indicate the median of each distribution (within-V1: 24 ms; within-V2: 74 ms). ‘***’: within-V2 GP timescales are significantly longer than within-V1 GP timescales (one-sided Wilcoxon rank sum test, *p* < 0.001). Bottom: V1a-V1b results. Normalized distribution of within-V1a (light blue) and within-V1b (dark blue) GP timescales across all 40 datasets (total within-V1a latent variables: 100; total within-V1b latent variables: 92). Triangles indicate the median of each distribution (within-V1a: 20 ms; within-V1b: 23 ms). ‘*’: within-V1b GP timescales are significantly longer than within-V1a GP timescales (one-sided Wilcoxon rank sum test, *p* < 0.05), even though the magnitude of the difference is small (as expected for randomly assigned subpopulations).

We further sought to characterize the strength—in addition to the dimensionality—of across-versus within-area activity in each area. We therefore considered the latent variables in V1 and in V2 separately, and computed the fraction of shared variance that each latent variable explained in its corresponding area (in Fig. 5, the amplitude of each latent time course is scaled by this value). Across-area variables explained only a portion of the shared variance in V1 and in V2 (Fig. 6b, top; median across-area strengths: 34% in V1; 76% in V2). Interestingly, across-area activity explained more of the shared variance in V2 than in V1 (Fig. 6b, top, points above the diagonal). This observation could not be fully attributed to differences in recorded population size or in the total dimensionality of each area (Supplementary Fig. 10). This discrepancy in across-area strength might be a consequence of the cortical layers from which we recorded: much of the activity in the middle layers of V2 is likely driven by V1. The superficial layers of V1, on the other hand, receive input from other sources that do not also project to the middle layers of V2.

Collectively, these observations (Fig. 6a,b, top) are consistent with the presence of a communication subspace between V1 and V2 [39], through which only a subset of population activity patterns are shared between the two areas. Our results further suggest that not only does there exist activity in V1 that is not shared with V2 (as reported in [39]), but there also exists activity in V2 that is not shared with V1. By contrast, V1a and V1b do not interact selectively. V1a-V1b “across-population” activity was consistently higher-dimensional (Fig. 6a, bottom; median dimensionality across populations 11; within-V1a: 2; within-V1b: 1), and accounted for nearly all of the shared variance in V1a and in V1b (Fig. 6b, bottom; median across-population strengths: 96% in V1a; 98% in V1b; note also the small amplitudes of the “within-population” latent time courses in Fig. 5b).

DLAG’s latent variables enabled further qualitative characterization of the moment-to-moment nature of within- and across-area activity on individual trials. For instance, stereotyped periodic signals, whose periods matched the period of the drifting grating presented, appeared strongly within V1 (Fig. 5a, top, “Across 3”, “Within 1”, and “Within 2”) and only weakly in V2 (Fig. 5a, bottom, “Across 3”). The prominence of this stimulus-related periodic structure in V1 relative to V2 is consistent with the stimulus response properties of neurons in each area [48], evident in the neuronal PSTHs (Fig. 4b). Care should be taken, however, when interpreting these latent variables as across-area interactions (see Discussion). By contrast, periodic signals were not evident in V1a or V1b within-population variables, but were evident in the activity shared between V1a and V1b (Fig. 5b, “Across 1” and “Across 2”). Other latent variables, particularly within V2, exhibited additional trial-to-trial variability whose connection to the presented stimulus is less apparent (for example, Fig. 5a, bottom, “Within 1” and “Within 2”).

#### V1-V2 interactions are bidirectional and asymmetric

We next used DLAG to study the bidirectional nature of interactions between V1 and V2. Each of DLAG’s across-area latent variables is associated with a time delay that indicates a feedforward (positive delay: V1 to V2) or feedback (negative delay: V2 to V1) interaction. For example, the first representative V1-V2 across-area variable (Fig. 5a, “Across 1”) was associated with a −23 ms delay, implying a feedback interaction. In contrast, the visually similar V1a-V1b across-population variable (Fig. 5b, “Across 3”) was associated with a 0 ms delay. A V1a-V1b delay at or near zero is expected, given that the V1a and V1b populations belong to the same area, and likely receive common inputs with similar latencies (in contrast to the populations in distinct areas V1 and V2).

We developed a statistical procedure to test whether such delays significantly deviate from zero. In brief, we assessed whether setting the delay to 0 ms resulted in a significant reduction in model performance; if so, the delay was deemed significant (i.e., “non-zero”; see Methods). Indeed, the directionality of this latent variable (“Across 3” for V1a-V1b) was identified as statistically “ambiguous” (i.e. not significantly different from zero, indicated by the bidirectional gray arrow in Fig. 5b).

Delays across all datasets reflected bidirectional interactions between V1 and V2 (Fig. 6c, top). Notably, the delays between V1 and V2 exhibited a striking asymmetry. The interactions across these areas were predominantly directed from V2 to V1 (Fig. 6c, top; median over “non-zero” delays: −8 ms; median over all delays: −5 ms). Among the across-area latent variables with statistically significant delays, 76% were associated with a negative delay. This asymmetry remained even when we subsampled the V1 population to match V2 in size, and re-applied DLAG (Supplementary Fig. 10). Like the strength of across-area activity observed in V1 and in V2 (Fig. 6b, top), the magnitudes of the delays might also reflect the cortical layers from which we recorded. The positive delays tended to be short (Fig. 6c, top; median across significant positive delays: +7 ms), consistent with the fact that the superficial layers of V1 directly project to the middle layers of V2 [25, 27]. The negative delays tended to be longer (Fig. 6c, top; median across significant negative delays: −11 ms), consistent with a multi-synaptic path from the middle layers of V2 back to the superficial layers of V1.

By contrast, V1a-V1b interactions were symmetric (Fig. 6c, bottom; median over “non-zero” delays: −2 ms; median over all delays: 0 ms; neither median significantly different from zero; 54% of “non-zero” delays were negative). This centering of the delay distribution around zero is expected, given that the neurons in V1a and V1b were randomly chosen and belong to the same area. Still, the magnitudes of V1a-V1b delays were not universally zero. These non-zero delays likely reflect aggregate differences in the stimulus response properties of the randomly chosen V1a and V1b subpopulations. For example, inspection of PSTHs (Fig. 4b) suggests that the phase of trial-averaged periodic structure can vary by tens of ms between individual V1 neurons.

Finally, we examined the timescales of neural activity identified by DLAG within V1 and V2. Within-V2 Gaussian process (GP) timescales were longer than within-V1 GP timescales (Fig. 6d, top; median within-V1: 24 ms; within-V2: 74 ms). Within-V1a and within-V1b GP timescales, on the other hand, were nearly the same (Fig. 6d, bottom; median within-V1a: 20 ms; within-V1b: 23 ms). These observations are consistent with previous evidence that timescales increase for areas higher up the cortical hierarchy [49, 50].

## Discussion

DLAG provides a novel description of population-level signal flow between populations of neurons. By leveraging the correlated activity across the two populations, DLAG can disentangle concurrent signals relayed in each direction and characterize how those signals evolve within and across trials. We demonstrated that DLAG performs well on synthetic datasets similar in scale to current neurophysiological recordings. Then we used DLAG to study bidirectional interactions between V1 and V2. Our framework lays a foundation for understanding how bidirectional signaling contributes to cortical function.

To our knowledge, DLAG has enabled for the first time the identification of bidirectional interactions between brain areas from spiking activity of neuronal populations. DLAG uncovered signatures of inter- and intra-areal interaction that are consistent with previous work, such as the selectivity with which V1 and V2 interact [39], as well as an increase in timescale moving up the cortical hierarchy from V1 to V2 [49, 50]. In addition, DLAG provided a novel ability to study the bidirectional nature of interactions between these areas, and characterize these interactions on a moment-to-moment basis. DLAG identified population-level interactions in both directions, whose strengths and associated time delays appear to reflect the cortical layers from which we recorded. Given our recording arrangement, we would expect DLAG to identify at least as many feedforward (V1 to V2) interactions as feedback (V2 to V1). Feedback connections do not originate in the input layers of V2; and, more generally, feedback inter-cortical connections equal feed-forward connections in number [46, 47], while feedback signals appear modulatory (rather than driving) in nature [51]. Surprisingly, DLAG revealed a marked asymmetry, such that a majority of across-area latent variables were associated with a feedback interaction. This apparent disparity presents an opportunity for future study.

Although we applied DLAG to the spiking activity of populations of neurons in distinct brain areas, DLAG is applicable to any high-dimensional time series data, including other neural recording modalities (e.g., calcium imaging). It can also be used to study the interaction of two populations of neurons in different cortical layers or of different cell types. DLAG can even be used to study the relationship between a neuronal population and a dynamic stimulus or behavioral variables.

### Relation to previous statistical methods

In contrast with other multivariate time series methods, such as Granger causal modeling [52–54], Generalized Linear Models [40, 55, 56], or recurrent neural networks [57], DLAG identifies low-dimensional across- and within-area latent variables with time delays and timescales. These latent variables enable a population-level characterization not only of activity that is shared across areas, but also of activity that is not. Existing time series methods that do incorporate dimensionality reduction may discard such within-area activity as noise [58, 59].

DLAG offers unique advantages when characterizing the temporal structure of activity within and across areas. Applied to V1 and V2, DLAG uncovered latent variables with diverse temporal profiles and timescales. The ability to capture diverse dynamical motifs stems from DLAG’s definition via Gaussian processes [60]: beyond temporal smoothness, DLAG makes no additional assumptions about the form of dynamics within or across areas. In contrast, multi-area methods proposed by [61] and [62], for instance, assume interactions evolve over time according to particular parametric (e.g., linear) dynamical models. Gaussian processes provide DLAG with another advantage [63]: the ability to discover wide-ranging delays with high precision. Existing multiarea methods (all of which, above, are defined in discrete-time) are limited to delays restricted to be integer multiples of the sampling period or spike count bin width of neural activity.

With the conceptual and statistical advantages described above, DLAG is a powerful tool for exploratory data analysis. For example, after performing a new experiment, one can use DLAG to generate data-driven hypotheses about plausible dynamical motifs within and across areas. Then, one can test these hypotheses using a dynamical system-based approach, for example, data-constrained recurrent networks [57].

### Interpretation of DLAG’s latent variables and time delays

One might interpret the population activity patterns represented by DLAG’s across-area variables as distinct “channels” with which two areas communicate [44]. As with any statistical method, however, interpretation of the features extracted by DLAG is subject to ambiguities, particularly when not all relevant brain areas and neurons are recorded [64]. An across-area latent variable, for instance, could reflect an interaction between areas A and B that is direct or indirect, mediated by a third (unobserved) area C. Similarly, a within-area latent variable could reflect activity internal to one area, or it could reflect inputs sent from unrecorded neurons to one area but not the other.

The sign and magnitude of DLAG’s time delays can, however, narrow the set of hypotheses consistent with the data. We might reasonably suspect, for example, that short positive (V1 to V2) delays identified by DLAG reflect direct interactions from the output layers of V1 to the input layers of V2 (the layers from which we recorded) [25, 27]. Larger negative (V2 to V1) delays might instead indicate indirect interactions, given that the path from the input layers of V2 to the output layers of V1 involves multiple synapses. Some across-area latent variables were associated with delays statistically indistinguishable from zero (i.e., “ambiguous”), and could reflect a third case: common input from an unobserved source. Future experimental interventions could further disambiguate these cases.

A phenomenon widely recognized by cross-correlation studies [22–27] is the presence of correlations across areas due simply to common stimulus drive, rather than an inter-areal interaction. For DLAG, these stimulus driven effects can appear as an across-area variable. The stereotyped periodic signals evident in V1-V2 across-area latent variables (Fig. 5a; “Across 3”) are a likely example. If desired, one could control for these effects with straightforward preprocessing steps, such as the subtraction of PSTHs from single-trial responses, thereby emphasizing trial-to-trial fluctuations correlated across areas [39].

Assumptions explicit in the DLAG model definition warrant additional care when interpreting estimated delays. First, DLAG treats time delays as constant parameters. However, interactions between areas might not be constant across different trial epochs or different experimental (e.g., stimulus) conditions. Thus, we interpret a delay as a summary of the dominant direction of interaction associated with a population activity pattern throughout the course of an experiment. Similarly, neurons within the same area can respond to a common input with different latencies (evident in, for example, Fig. 4b). An estimated delay hence also represents a summary across neurons [63]. Moreover, DLAG assumes that each dimension of population activity is associated with one delay, or direction. If a set of interactions were to occur currently in both directions but evolve along the same dimension, then teasing apart directionality might be difficult—albeit for any statistical method, not just DLAG. Finally, DLAG assumes that signals are at most linearly transformed across areas. DLAG therefore does not take into account non-linear transformations of signals. We believe that there are many experimental scenarios for which the assumption of a linear transformation or direct signal transmission is appropriate (e.g., [15]). Nonetheless, in practice, this assumption should be evaluated on a case-by-case basis.

Solutions to these interpretational challenges might already be well within reach, if not already available through DLAG’s existing machinery. For example, one could fit DLAG to subsets of trials, subsets of neurons, or to separate trial epochs to understand how DLAG’s estimates depend on these elements of the neural recordings. We have already employed some of these strategies here (Fig. 6, Supplementary Fig. 10: our nonparametric bootstrap procedure for delay significance and our analyses of V1 subpopulations), and could continue to build upon that foundation.

## Methods

### Mathematical notation

To disambiguate each variable or parameter in the DLAG model, we need to keep track of up to four labels that indicate their associated (1) subpopulation (e.g., brain area); (2) neuron or latent variable index; (3) time point; or (4) designation as within- or across-area. We indicate the first three labels via subscripts, where subpopulations (areas) are indexed by *i* = 1,2; neurons or latent variables are indexed by *j* (we’ll indicate the upper bound as appropriate); and time is indexed by *t* = 1,…, *T*. For example, we define the observed activity of neuron *j* (out of *q_i_*) in area *i* at time *t* as 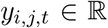. To indicate a collection of all variables along a particular index, we replace that index with the ‘:’ symbol. Hence we represent the simultaneous activity of the population of *q_i_* neurons observed in area *i* at time *t* as the vector 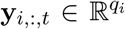. For concision, where a particular index is either not applicable or not immediately relevant, we omit it. The identities of the remaining indices should be clear from context. For example, throughout this work we consider only the activity of a full population, and not of single neurons, so we rewrite y_*i*_,:,*t* as y_*i*,*t*_. Finally, we indicate a latent variable’s or parameter’s designation as within- or across-area via a superscript, where ‘*w*’ indicates within-area, and ‘*a*’ indicates across-area. For example, we define across-area latent variable *j* (out of *p^a^*) in area *i* at time *t* as 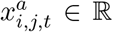, and the collection of all *p^a^* latent variables as the vector 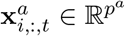. We similarly define within-area latent variable *j* (out of 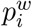) in area *i* at time *t* as 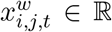, and the collection of all 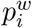 latent variables as the vector 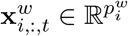.

It is conceptually helpful to understand the above notation for observed (y) and latent (x) variables as taking cross-sections of matrices. For example, observed activity in area *i* can be grouped into the matrix 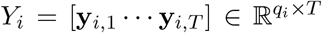. Then, each y_*i*,*t*_ is a column of *Y_i_*. Similarly, across-area latent variables in area *i* can be grouped into the matrix 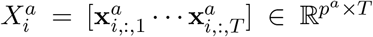. Each 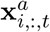 is a column of 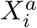. Similarly, we represent a row of 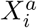 (i.e., the values of a single latent variable *j* at all time points) as 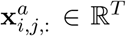. Within-area latent variables can be understood analogously from the matrix 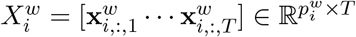.

We will explicitly define all other variables and parameters as they appear, but for reference, we list common variables and parameters below:

#### Observed neural activity

- *q_i_* – number of neurons observed in area *i*
- *Y_i_* – *q_i_* × *T* matrix of observed activity in area *i*
- *y*_*i*,*t*_ – *q_i_* × 1 vector of observed activity in area *i* at time *t*; the *t*^th^ column of *Y_i_*

#### Latent variables

- *p^a^* – number of across-area variables (same for both areas)
- 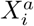 – *p^a^* × *T* matrix of across-area variables in area *i*
- 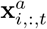 – *p^a^* × 1 vector of across-area variables in area *i* at time *t*; the *t*^th^ column of 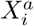
- 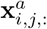 – *T* × 1 vector of values of across-area variable *j* in area *i* over time; the *j*^th^ row of 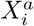
- 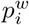 – number of within-area variables in area *i*
- 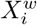 – 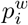 matrix of within-area variables in area *i*
- 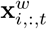 – 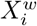 vector of within-area variables in area *i* at time *t*; the *t*^th^ column of 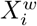
- 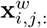 – *T* × 1 vector of values of within-area variable *j* in area *i* over time; the *j*^th^ row of 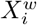

#### Model parameters

- 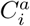 – *q_i_* × *p^a^* across-area loading matrix for area *i*
- 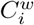 – 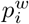 within-area loading matrix for area *i*
- d_*i*_ – *q_i_* × 1 mean parameter for area *i*
- *R_i_* – *q_i_* × *q_i_* observation noise covariance matrix for area *i*
- *D*_*i*,*j*_ – time delay parameter between area *i* and across-area variable *j*
- *D_j_* – relative time delay associated with across-area variable *j*; *D_j_* = *D*_2,*j*_ – *D*_1,*j*_
- 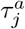 – Gaussian process timescale for across-area variable *j*
- 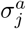 – Gaussian process noise parameter for across-area variable *j*
- 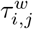 – Gaussian process timescale for within-area variable *j* in area *i*
- 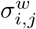 – Gaussian process noise parameter for within-area variable *j* in area *i*

#### Gaussian process covariances

- 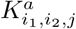 – *T* × *T* covariance matrix for across-area variable *j*, between areas *i*_1_ and *i*_2_
- 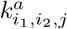 – covariance function for across-area variable *j*, between areas *i*_1_ and *i*_2_
- 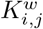 – *T* × *T* covariance matrix for within-area variable *j* in area *i*
- 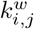 – covariance function for within-area variable *j* in area *i*

#### DLAG observation model

For area *i* at time *t*, we define a linear-Gaussian relationship between observed activity, y_*i*,*t*_, and latent variables, 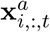 and 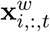 [65]:

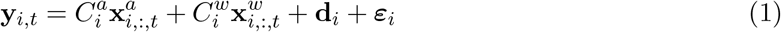

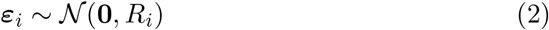

where 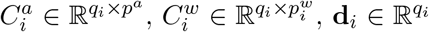, and 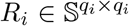 (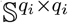 is the set of *q_i_* × *q_i_* symmetric matrices) are model parameters to be estimated from data. The relationship between observed and latent variables is illustrated graphically in Supplementary Fig. 1. The loading matrices 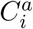 and 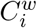 linearly combine latent variables and map them to observed neural activity. The parameter *d_i_* can be thought of as the mean firing rate of each neuron. *ε_i_* is a zero-mean Gaussian random variable, where we constrain the covariance matrix *R_i_* to be diagonal, as in factor analysis (FA) [45] and Gaussian process factor analysis (GPFA) [60], to capture variance that is independent to each neuron. This constraint encourages the latent variables to explain as much of the shared variance among neurons as possible.

As we will describe, at time point *t*, across-area variables 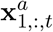 and 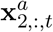 in area 1 and area 2, respectively, are coupled with each other, and thus each area has the same number of across-area variables, *p^a^*. Within-area variables are not coupled across areas, on the other hand, and thus each area *i* may have a different number of within-area variables, 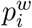. Because we seek a low-dimensional description of neural activity in each area, the combined number of across- and within-area variables is less than the number of neurons, i.e., 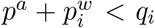, where *p^a^* and 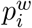 are determined by the data (see below).

The parameters 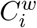 and 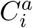 have an intuitive geometric interpretation (Fig. 2, right column). Each element of y_*i*,*t*_, the activity of each neuron in area *i*, can be represented as an axis in a high-dimensional population activity space. Then the columns of 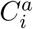, the across-area loading matrix for area *i*, define a subspace in this population activity space, where each dimension corresponds to a distinct across-area latent variable. This across-area subspace represents patterns of population activity that is correlated across areas. Analogously, the columns of 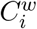 define a within-area subspace, which represents patterns of population activity that is shared only among neurons within area *i*. Additionally, as we will discuss below, since the *j*^th^ pair of across-area variables 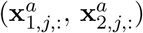 is associated with a direction of population signal flow (Fig. 2, center column), so too are the corresponding columns in 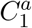 and 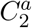. The across-area subspace can thus be partitioned further based on the nominal directionality of activity patterns (area 1 to area 2, or area 2 to area 1). Finally, note that the columns of 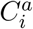 and 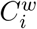 (and the subspaces they define) are linearly independent; but they are not, in general, orthogonal. The ordering of these columns, and of the corresponding latent variables, is arbitrary.

### DLAG state model

We seek to extract smooth, single-trial latent time courses, where the degree of smoothing is determined by the neural activity (as described below). The time course of each within-area and across-area latent variable is described by a Gaussian process (GP) [66].

#### Within-area latent variables

For each within-area variable 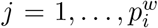 in brain area *i*, we define a separate GP as follows [60]:

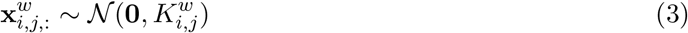

where 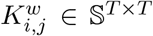 is the covariance matrix for within-area variable *j* of area *i*. DLAG is compatible with any valid form of GP covariance, but for the present work, we choose the commonly used squared exponential (SE) function. Then, element (*t*_1_,*t*_2_) of 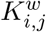, the covariance between samples of the within-area variable at times *t*_1_ and *t*_2_, can be computed according to:

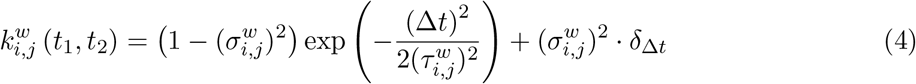

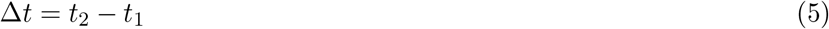

where the characteristic timescale, 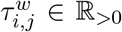, and GP noise variance, 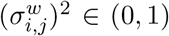, are model parameters. *δ*_Δ*t*_ is the kronecker delta, which is 1 for Δ*t* = 0 (equivalently, *t*_1_ = *t*_2_) and 0 otherwise.

Notice that 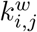 is stationary: the SE function depends only on the time difference (*t*_2_ – *t*_1_) (Supplementary Fig. 2a). This stationarity gives the covariance matrix 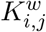 a characteristic banded structure (Supplementary Fig. 2b). The characteristic timescale, 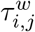, dictates the width of 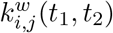, or equivalently, how rapidly the latent variable changes over time. The 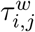 parameters are estimated from the neural activity, together with the other DLAG parameters (see below). We follow the same conventions as in [60], and fix 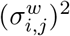 to a small value (10^-3^). Note also that, under this definition, the process is normalized so that 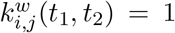 for *t*_1_ = *t*_2_. Thus, the prior distribution of within-area latent variables 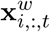 in area *i* at each time *t* follows the standard normal distribution, 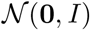. This normalization removes model redundancy in the scaling of 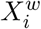 and 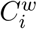.

Beyond describing within-area interactions, within-area variables are critical to the interpretability of across-area variables. As we will define below, across-area variables describe the activity of neurons in both areas. Within-area variables could, in principle, be formulated as a special case of across-area variables, where the loading coefficients to one area (the appropriate columns of 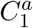 or 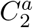 in equation (1)) are identically zero. If the model does not allow for within-area variables, then across-area variables must explain within-area activity in addition to across-area activity. Across-area variables could thus reflect a mixture of within- and across-area activity in this case, obfuscating their interpretation as representing population activity patterns that are correlated across areas. The presence of within-area variables allows the across-area variables to isolate activity that is truly correlated across areas. This statistical phenomenon applies to other statistical models, and is not specific to DLAG [34, 61]. See Supplementary Discussion for further mathematical discussion.

#### Across-area latent variables

We next describe across-area temporal structure. Across-area variables are different from within-area variables in two respects: (1) across-area variables are defined in pairs, where the elements of each pair correspond to the two areas, and (2) the elements of each pair are time-delayed relative to each other (Fig. 2, center column). Thus in contrast to our definition of within-area variables, in which we considered each area separately, we now consider across-area variables in both areas together: 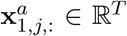 and 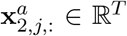, the *j*^th^ rows of 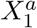 and 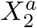, respectively, for the *j*^th^ across-area variable.

The across-area latent variables of area 1 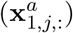 and area 2 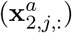 belong to the same GP (Supplementary Fig. 2c). The 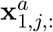 are values of the GP sampled on a time grid. The 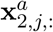 are values of the same GP, also sampled on a time grid, but offset from the time grid of area 1 by a time delay. We define the GP for each across-area variable *j* = 1,…, *p^a^* as follows:

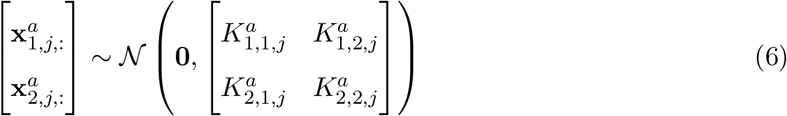

where 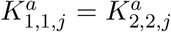 describe the autocovariance of each across-area variable, and 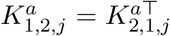 describe the cross-covariance that couples the two areas (Supplementary Fig. 2d).

To express the auto- and cross-covariance functions, we introduce additional notation. Specifically, we indicate brain areas with two subscripts, *i*_1_ = 1, 2 and *i*_2_ = 1, 2. Then, we define 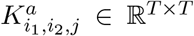 to be either the auto- or cross-covariance matrix between across-area variable 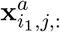 in area *i*_1_ and across-area variable 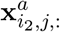 in area *i*_2_. We again choose to use the SE function for GP covariances. Therefore, element (*t*_1_,*t*_2_) of each 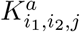 can be computed as follows [63]:

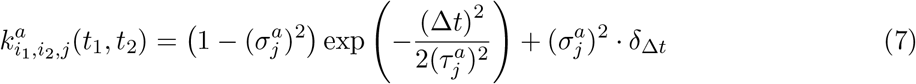

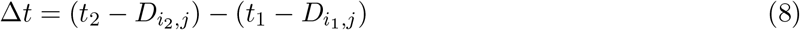

where the characteristic timescale, 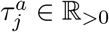, and the GP noise variance, 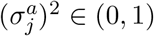, are model parameters. *δ*_Δ*t*_ is the kronecker delta, which is 1 for Δ*t* = 0 and 0 otherwise.

We also introduce two new parameters: the time delay to area *i*_1_, 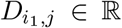, and the time delay to area *i*_2_, 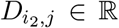. Notice that, when computing the autocovariance for area *i* (i.e., *i*_1_ = *i*_2_ = *i*), the time delay parameters *D*_*i*1,*j*_ and *D*_*i*2,*j*_ are equal, and so Δ*t* (equation (8)) reduces simply to the time difference (*t*_2_ – *t*_1_), as in the within-area case (equation (5)). Time delays are therefore only relevant when computing the cross-covariance between area 1 and area 2. The time delay to area 1, *D*_1,*j*_, and the time delay to area 2, *D*_2,*j*_, by themselves have no physically meaningful interpretation. Their difference *D_j_* = *D*_2,*j*_ – *D*_1,*j*_, however, represents a well-defined, continuous-valued time delay from area 1 to area 2. The sign of the relative time delay *D_j_* indicates the directionality of the lead-lag relationship between areas captured by latent variable *j* (positive: area 1 leads area 2; negative: area 2 leads area 1), which we interpret as a description of inter-areal signal flow.

Both the characteristic timescales 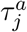 and relative delays *D_j_* are estimated from the neural activity, together with the other DLAG parameters (see below). More specifically, to ensure identifiability of time delay parameters, we designate area 1 as the reference area, and fix the delays for area 1 at 0, that is, *D*_1,*j*_ = 0 for all across-area variables *j* = 1,…, *p^a^*. Then, each relative time delay *D_j_* is simply the time delay parameter to area 2, *D*_2,*j*_. Note that *D_j_* need not be an integer multiple of the sampling period or spike count bin width of the neural activity. As in the within-area case, the across-area GP noise variance, 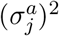, is set to a small value (10^-3^). Furthermore, the across-area GP is also normalized so that 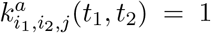 if Δ*t* = 0, thereby removing model redundancy in the scaling of 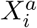 and 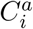.

#### DLAG special cases

Finally, we consider some special cases of the DLAG model that illustrate its relationship to other dimensionality reduction methods. First, by fixing all time delays to zero (*D_j_* = 0), and by removing within-area latent variables 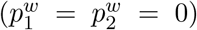, DLAG becomes equivalent to Gaussian process factor analysis (GPFA) [60] applied to both areas jointly. By removing instead the across-area latent variables (*p^a^* = 0), and keeping the within-area latent variables intact, DLAG becomes equivalent to GPFA applied to each area independently. And finally, by removing temporal smoothing (i.e., in the limit as all GP noise parameters 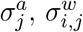 approach 1), while keeping both within- and across-area latent variables, DLAG becomes similar to probabilistic canonical correlation analysis (pCCA) [65, 67]. Whereas pCCA describes within-area activity via observation noise covariance matrices (*R_i_*; see equation (37)), this special-case DLAG model would describe within-area activity via low-dimensional latent variables.

### Fitting the DLAG model

Equations (1)–(8) provide a full definition of the DLAG model. In this section, we describe how DLAG model parameters are fit using exact Expectation Maximization (EM), where the parameters are

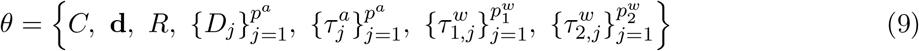

Toward that end, we first write the DLAG observation model more compactly as follows. Define the joint activity of neurons in all brain areas by vertically concatenating the observations in each area, y_1,*t*_ and y_2,*t*_:

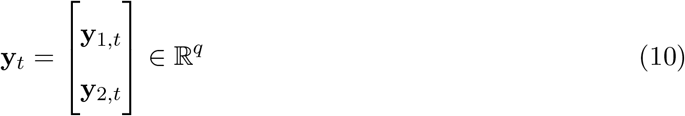

where *q* = *q*_1_ + *q*_2_. Next we group together the across- and within-area latent variables for the *i*^th^ brain area to define 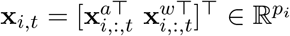, where 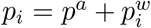. We then vertically concatenate the latent variables in each area:

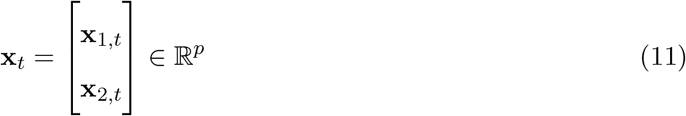

where *p* = *p*_1_ + *p*_2_. We also define the following structured matrices. First define 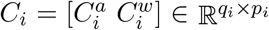 by horizontally concatenating 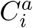 and 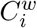. Then, we collect the *C_i_* into a block-diagonal matrix as follows:

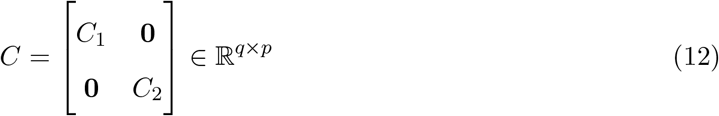

Similarly, define

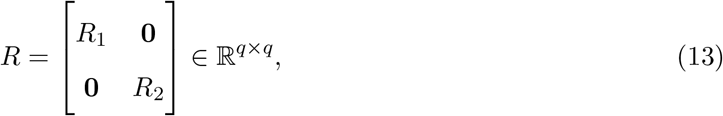

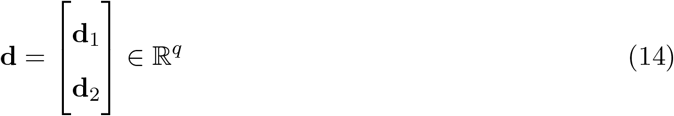

We can then write the DLAG observation model compactly as follows:

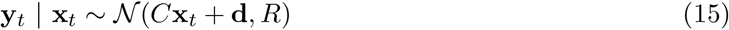

The observation model expressed in equation (15) defines a distribution for neural activity at a single time point, but to properly fit the DLAG model, we must consider the distribution over all time points. Thus we define 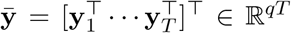 and 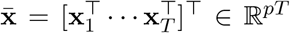, obtained by vertically concatenating the observed variables y_*t*_ and latent variables x_*t*_, respectively, across all *t* = 1,…, *T*. Then, we rewrite the state and observation models as follows:

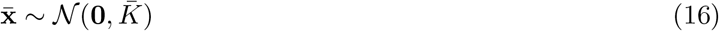

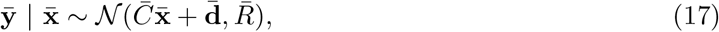

where 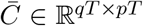 and 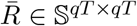 are block diagonal matrices comprising *T* copies of the matrices *C* and *R*, respectively. 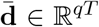 is constructed by vertically concatenating *T* copies of d. The elements of 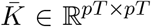 are computed using equations (3)–(8). Then, the joint distribution over observed and latent variables is given by

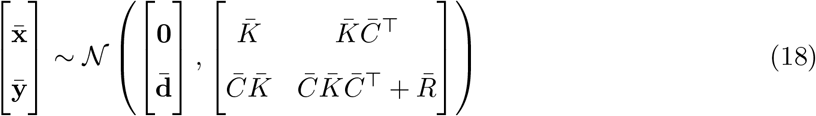

#### E-step

In the E-step, our goal is to compute the posterior distribution of the latent variables 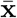 given the recorded neural activity ȳ, 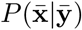, using the most recent parameter estimates *θ*. Using basic results of conditioning for jointly Gaussian random variables, we get

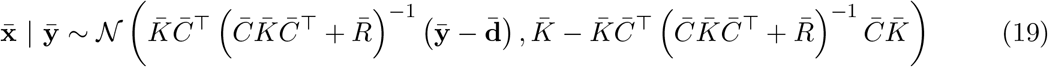

Thus, posterior estimates of latent variables are given by

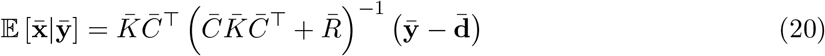

The marginal likelihood of the observed neural activity can be computed as

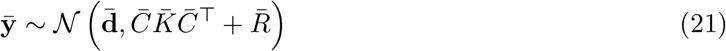

#### M-step

In the M-step, our goal is to maximize 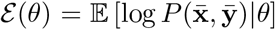 with respect to *θ*, using the latest inference of the latent variables, computed in the E-step. As in [60, 63], we adopt the following notation. Given a vector v,

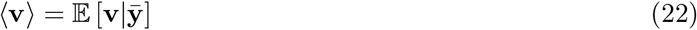

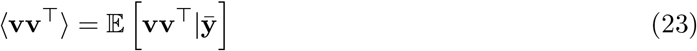

The appropriate expectations can be found using equation (19).

Maximizing *ε*(*θ*) with respect to *C*, d yields the following closed-form update for the *i*^th^ brain area:

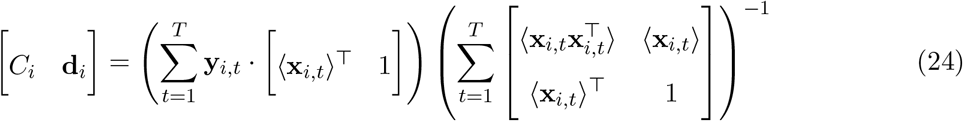

After performing the update for each area separately, we collect all updated values into *C* and d.

Then we update *R* for both brain areas together, as follows:

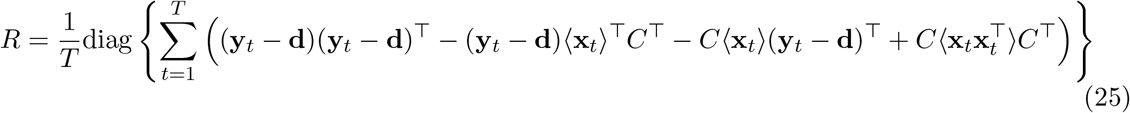

There are no closed-form solutions for the Gaussian process parameter updates, but we can compute gradients and perform gradient ascent. Note that, for this work, we choose not to fit the Gaussian process noise variances, but rather, we set them to small values (10^-3^), as in [60]. Within-area timescale gradients for the *i*^th^ brain area and *j*^th^ within-area latent variable are given by

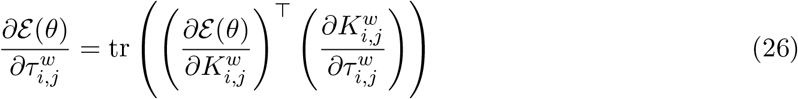

where

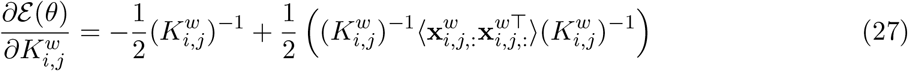

and element (*t*_1_, *t*_2_) of 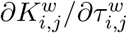 is given by

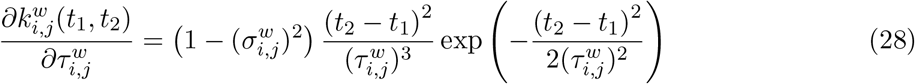

To express the across-area timescale and delay parameter gradients, we introduce more compact notation for the variables in equation (6). Let 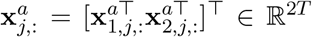 for the *j*^th^ across-area latent variable, and

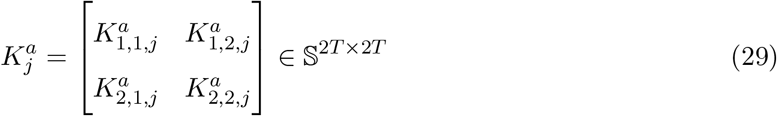

Then, across-area timescale gradients are given by

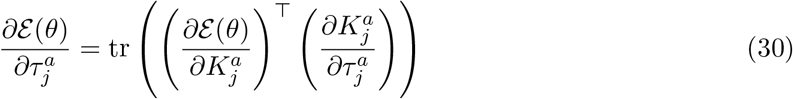

where

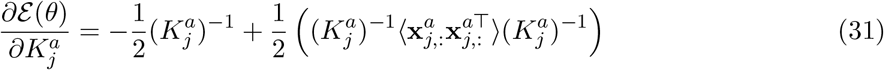

and each element of 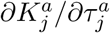 is given by

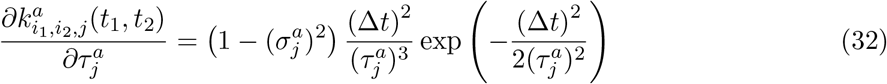

where Δ*t* is defined as in equation (8). To optimize the timescales while respecting non-negativity constraints, we perform a change of variables, and then perform unconstrained gradient ascent with respect to 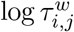 or 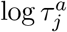.

Next, delay gradients for brain area *i* and across-area latent variable *j* are given by

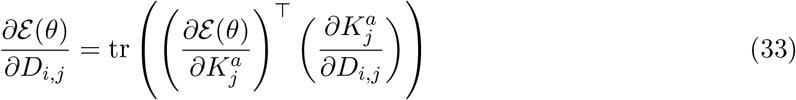

where 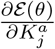 is defined as in equation (31), and each element of 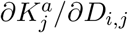 is given by

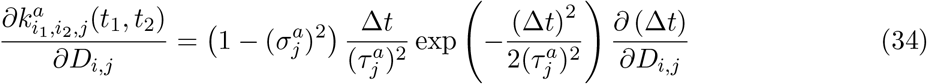

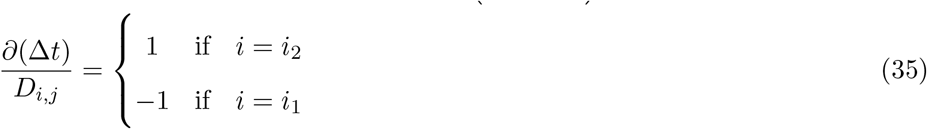

where Δ*t*, *i*_1_, and *i*_2_ are defined as in equation (8). In practice, we fix all delay parameters for area 1 at 0 to ensure identifiability. As with the timescales, one might wish to constrain the delays within some physically realistic range, such as the length of an experimental trial, so that −*D*_max_ ≤ *D_i,j_* ≤ *D*_max_. Toward that end, we make the change of variables 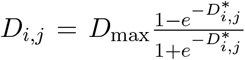 and perform unconstrained gradient ascent with respect to 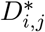. Here we chose *D*_max_ to be half the length of a trial. No delays came close to these constraints in our results (Fig. 6, Supplementary Fig. 10).

Finally, note that all of these EM updates are derived for a single sequence, or trial. It is straight-forward to extend these equations to *N* independent sequences (each with a potentially different number of time steps, *T*) by maximizing 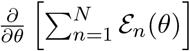.

#### Parameter initialization

To initialize the DLAG observation model parameters to reasonable values prior to fitting with the EM algorithm, we first fit a probabilistic canonical correlation analysis (pCCA) [67] model to the neural activity, with the same number of across-area latent variables as the desired DLAG model (see next section). pCCA is defined by the following state and observation models:

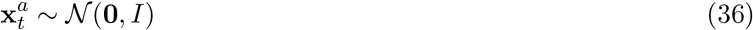

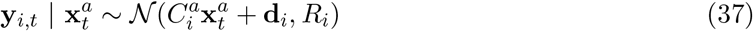

where 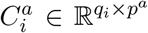 maps the *p^a^*-dimensional across-area latent variables 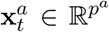 to the neural activity of area *i*, 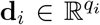 is a mean parameter, and 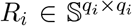 is the observation noise covariance matrix. *R_i_* is not constrained to be diagonal. The fitted values for 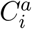 and d_*i*_ are used as initial values for their DLAG analogues. We take only the diagonal elements of *R_i_* to initialize its DLAG analogue.

pCCA does not incorporate within-area latent variables. Therefore, we initialized each DLAG within-area loading matrix 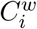 so that its columns spanned a subspace uncorrelated with that spanned by the columns of 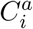, returned by pCCA. Such a subspace can be computed as follows. Let 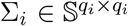 be the sample covariance matrix of activity in area *i*. Then define 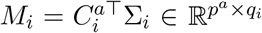. The singular value decomposition of *M_i_* is given by 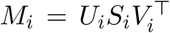, where 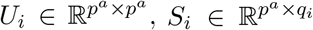, and 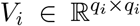. The first *p^a^* columns of *V_i_* span the same across-area subspace spanned by the columns of 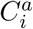. The remaining *q_i_* – *p^a^* columns form an orthonormal basis for the subspace uncorrelated with this across-area subspace. We initialized 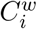 with the first 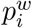 of these uncorrelated basis vectors. Finally, we initialized all delays to zero, and all within- and across-area Gaussian process timescales to the same value, equal to twice the sampling period or spike count bin width of the neural activity.

### Selecting the number of within- and across-area latent variables

DLAG has three hyperparameters: *p^a^*, the number of across-area latent variables; and 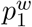 and 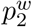, the number of within-area latent variables for each area. Model selection therefore poses a significant scaling challenge. Grid search over even a small range of within- and across-area dimensionalities can result in a large number of models that need to be fitted and validated. For example, considering just 10 possibilities for each type of latent variable would result in 1,000 model candidates. Thus, exhaustive search for the optimal DLAG model is impractical.

We therefore developed a streamlined cross-validation procedure that significantly improves scalability. In brief, our model selection procedure occurs in two stages. First, we consider each area separately, and—using factor analysis (FA) [45]—we find the number of latent variables needed to explain the shared variance among neurons within each area. We reasoned that, while there is not a direct correspondence between the optimal number of latent variables in DLAG and FA models (because of temporal smoothing and other differences in model structure), it is unlikely that the total number of within- and across-area latent variables extracted by DLAG will exceed the FA dimensionality for an area (such a case would imply that there exists a neuron in, for example, area A that covaries with one or more neurons in area B, but no other neurons in area A). Hence we believe this approach to be reasonable given the significant computational benefits. We then use the FA dimensionality in each area to reduce the space of DLAG model candidates to a practical size.

In greater detail, we first applied FA to each area independently, and identified the optimal FA dimensionality through *K*-fold cross-validation (here we chose *K* = 4). The FA model with the highest cross-validated data likelihood was taken as “optimal.” We then used the optimal FA dimensionalities 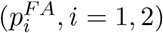 to constrain the space of DLAG model candidates. In particular, we consider only DLAG models that satisfy 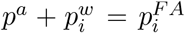, for *i* = 1, 2; and 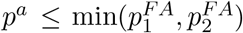. In words, we consider only DLAG models such that the number of within- and across-area latent variables in each area sum to that area’s optimal FA dimensionality. Furthermore, the number of across-area latent variables is limited by the area with the smallest optimal FA dimensionality.

Not only does this streamlined cross-validation approach provide an upper limit on the possible number of within- and across-area latent variables, it also effectively collapses the DLAG hyperparameter space from three free hyperparameters to one (across-area dimensionality, *p^a^*), drastically improving scalability. Among the model candidates within this constrained search range, we selected models that exhibited the largest cross-validated data likelihood (computed as in equation (21)), using the same *K*-fold cross-validation scheme as for FA. To further reduce runtime, we limited the number of EM iterations during cross-validation to 1,000. The optimal DLAG model was then re-fit to full convergence, where the data log-likelihood improved from one iteration to the next by less than a preset tolerance (here we used 10^-8^).

We also note that throughout this work, we explicitly considered model candidates for which across-area dimensionality was zero (*p^a^* = 0): the two areas are independent, and any correlations between neurons are purely within-area. Similarly, we explicitly considered model candidates for which within-area dimensionalities were zero (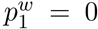 or 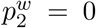): all variance shared among neurons in one area is attributed to their interactions with neurons in the other area. The case where all dimensionalities are zero 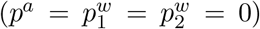 is equivalent to fitting a multivariate Gaussian distribution to the data with diagonal covariance (i.e., all neurons are treated as independent). We similarly considered zero-dimensional FA models (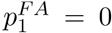 or 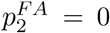) during the first stage of our model selection procedure, equivalent to fitting a multivariate Gaussian distribution with diagonal covariance to observations in the respective area. The inclusion of these zero-dimensionality model candidates protects against the identification of spurious interactions across or within areas.

### Synthetic data generation

We generated synthetic datasets according to the DLAG generative model, so that we could leverage known ground truth to evaluate the accuracy of estimates and characterize DLAG’s performance over a range of simulated conditions. We started by randomly generating the set of model parameters, *θ*, subject to constraints informed by experimental data. For all datasets, we chose the numbers of neurons in each area based on our V1-V2 recordings (area A: *q*_1_ = 80; area B: *q*_2_ = 20). We set the combined total dimensionality in each area to representative values (area A: 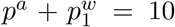; area B: 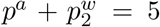), but varied the relative number of within- and across-area latent variables across datasets. Generating 20 datasets at each of six configurations (*p^a^* = 0,…, 5; 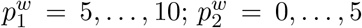) resulted in a total of 120 independent datasets. Importantly, among these datasets, we included datasets without across- or within-area structure (i.e., datasets for which across- or within-area dimensionality was zero), to test if our framework could identify such cases.

To ensure that synthetic datasets exhibited realistic noise levels, we first evaluated the strength of latent variables relative to the strength of single-neuron variability exhibited in the V1-V2 recordings. Specifically, we computed the “signal-to-noise” ratio (where “signal” is defined as the shared activity described by latent variables), 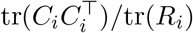, for V1 and V2 using the parameters of the optimal DLAG models fit to each V1-V2 dataset. Representative values were 0.3 and 0.2 for V1 and V2, respectively. Then for each dataset, we generated our synthetic observation model parameters, *C^i^* and *R_i_*, as follows. We first drew the elements of *C_i_* and a diagonal matrix 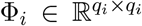 from the standard normal distribution 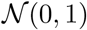. Then, we set 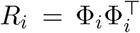 (so that *R_i_* was a valid covariance matrix) and rescaled *R_i_* such that area i exhibited the correct signal-to-noise ratio. The elements of the mean parameter d were also drawn from the standard normal distribution.

Finally, we drew all timescales 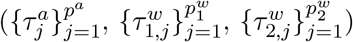 uniformly from *U*(*τ*_min_, *τ*_max_), with *τ*_min_ = 10 ms and *τ*_max_ = 150 ms. We drew all delays ({*D*_1_,…, *D_p^a^_*}) uniformly from *U*(*D*_min_, *D*_max_), with *D*_min_ = –30 ms and *D*_max_ = +30 ms. All Gaussian process noise variances 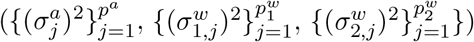 were fixed at 10^-3^. With all model parameters specified, we then generated *N* = 100 independent and identically distributed trials 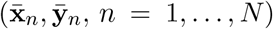 according to equations (16) and (17). Each trial comprised *T* = 50 time points, corresponding to 1, 000 ms sequences sampled with a period of 20 ms, to mimick the 20 ms spike count time bins used to analyze the experimental data.

### Synthetic data performance metrics

To quantify DLAG’s performance across all synthetic datasets, we employed a variety of metrics. We first consider the estimation of DLAG’s observation model parameters. To assess the accuracy of loading matrix estimation (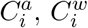; reported in Fig. 3, Supplementary Fig. 3, and Supplementary Fig. 4), we computed a normalized subspace error [68]:

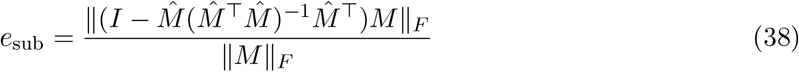

where *M* is the appropriate ground truth parameter, 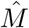 is the corresponding estimate, and ||·||_*F*_ is the Frobenius norm. *e*_sub_ quantifies the magnitude of the projection of the column space of *M* onto the null space of 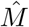. A value of 1 indicates that the column space of *M* lies completely in the null space of 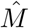, and therefore the estimate captures no component of the ground truth. A value of 0 indicates that the column space of 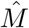 contains the full column space of *M*, and therefore the estimate captures all components of the ground truth. This metric offers two advantages: (1) it does not require that the columns of *M* and 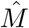 are ordered in any way (the ordering of DLAG latent variables is arbitrary); and (2) it does not require that *M* and 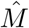 have the same number of columns, so it can be used to compare the performance of models with different numbers of latent variables. We report the accuracy of loading matrix estimation as 1 – *e*_sub_ (Fig. 3). To assess the accuracy of estimating d and *R* (reported in Supplementary Fig. 3 and Supplementary Fig. 4), we computed the normalized error

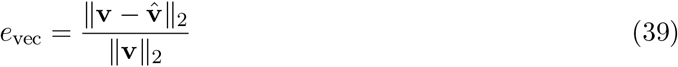

where v is either d or diag(*R*), and 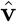 is the corresponding estimate.

We next consider the estimation of DLAG’s state model parameters. Reporting the accuracy of delay and timescale estimates (Fig. 3, Supplementary Fig. 3, Supplementary Fig. 4, and Supplementary Fig. 5) required explicitly matching estimated latent variables to the ground truth. Given the large number of synthetic datasets presented here, we automated this matching process as follows. First, for each area i, we took the unordered across- and within-area latent variable estimates, 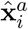 and 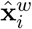, and computed the pairwise correlation between each estimated latent variable and each ground truth latent variable, 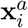 and 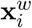, across all time points and trials. We then reordered the estimated latent variables to match the ground truth latent variables with which they showed the highest magnitude of correlation. To report delay and timescale estimation performance, we computed the absolute error between ground truth and (matched) estimated parameters, to express the error in units of time (ms).

Finally, we consider the moment-by-moment estimation of latent variables. As with the loading matrix, delay, and timescale estimates, quantifying the accuracy of latent variable estimates requires care since the sign and ordering of latent variables is arbitrary and will not, in general, match between estimates and the ground truth. First, let 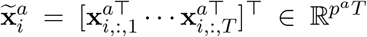 be a collection of all (ground truth) across-area variables at all time points in area *i*. Similarly, let 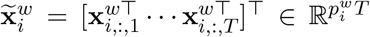 be a collection of all (ground truth) within-area variables at all time points in area *i*. Finally, define 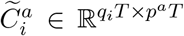 and 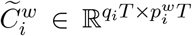 to be block diagonal matrices comprising *T* copies of the (ground truth) matrices 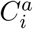 and 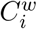, respectively; and define 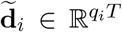 by vertically concatenating *T* copies of (the ground truth) d_*i*_. We’ll denote the estimates of each of these values by 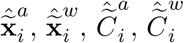, and 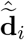. The estimates 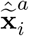 and 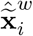 are posterior means, computed according to equation (20).

Then, to separate the accuracy of across-area variable estimation from the accuracy of within-area variable estimation (as reported in Fig. 3, Supplementary Fig. 3, and Supplementary Fig. 4), we estimated denoised (smoothed) observations, using only across-area or only within-area latent variable estimates:

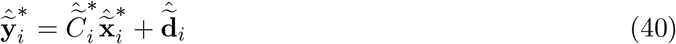

where 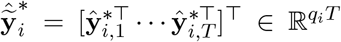. Here, the ‘*’ symbol is used to indicate either *a* or *w* as a superscript, where observations have been denoised using only across- or within-area variable estimates, respectively. We then collect the denoised sequences on all *N* trials, 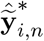, *n* = 1,…, *N*, into the matrix 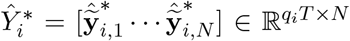. Analogously, define 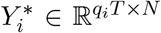 to be the set of ground truth sequences generated prior to adding noise (i.e., the noise term *ε_i_*, defined in equation (2)).

We then computed the *R*^2^ value between estimated and (noiseless) ground truth sequences:

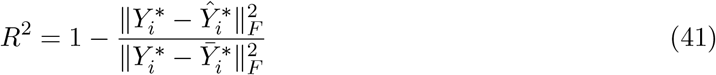

where 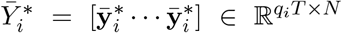 is constructed by horizontally concatenating *N* copies of the sample mean for each neuron in the ground truth 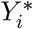, taken over all time points and trials 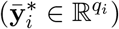. Note that, in the multivariate case, *R*^2^ ∈ (–∞, 1], where a negative value implies that estimates predict the ground truth less accurately than simply the sample mean.

### Visual stimuli and neural recordings

Animal procedures and recording details have been described in previous work [27, 69]. Briefly, animals (macaca fascicularis, young adult males) were anesthetized with ketamine (10 mg/kg) and maintained on isoflurane (1%-2%) during surgery. Recordings were performed under sufentanil (typically 6-18 mg/kg/hr) anesthesia. Vecuronium bromide (150 mg/kg/hr) was used to prevent eye movements. The duration of each experiment (which comprised multiple recording sessions) varied from 5 to 7 days. All procedures were approved by the IACUC of the Albert Einstein College of Medicine.

The data analyzed here are those reported in [39, 44], and a subset of recording sessions reported in [27]. Activity in V1 output layers was recorded using a 96 channel Utah array (400 micron inter-electrode spacing, 1 mm length, inserted to a nominal depth of 600 microns; Blackrock, UT). We recorded V2 activity using a set of electrodes/tetrodes (interelectrode spacing 300 microns) whose depth could be controlled independently (Thomas Recording, Germany). These electrodes were lowered through V1, the underlying white matter, and then into V2. Within V2, we targeted neurons in the input layers. We verified the recordings were performed in the input layers using measurements of the depth in V2 cortex, histological confirmation (in a subset of recordings), and correlation measurements. For complete details see [69] and [27]. Voltage snippets that exceeded a user-defined threshold were digitized and sorted offline. The sampled neurons had spatial receptive fields within 2-4° of the fovea, in the lower visual field.

We measured responses evoked by drifting sinusoidal gratings (1-1.1 cyc/°; drift rate of 6.25 Hz; 2.6-4.95° in diameter; full contrast, defined as Michelson contrast, (*L*_max_ – *L*_min_)/(*L*_max_ + *L*_min_), where *L*_min_ is 0 cd/m^2^ and L_max_ is 80 cd/m^2^) at 8 different orientations (22.5° steps), on a calibrated CRT monitor placed 110 cm from the animal (1024 x 768 pixel resolution at a 100 Hz refresh rate; Expo: http://sites.google.com/a/nyu.edu/expo). Each stimulus was presented 400 times for 1.28 seconds. Each presentation was preceded by an interstimulus interval of 1.5 seconds during which a gray screen was presented.

We recorded neuronal activity in three animals. In two of the animals, we recorded in two different but nearby locations in V2, providing distinct middle-layer populations, yielding a total of five recording sessions. We treated responses to each of the 8 stimuli in each session separately, yielding a total of 40 “datasets.”

### Data preprocessing

We counted spikes in 20 ms time bins during the 1.28 second stimulus presentation period (64 bins per trial). For all analyses corresponding to each recording session, we excluded neurons that fired fewer than 0.5 spikes/second, on average, across all trials and all grating orientations. Because we were interested in V1-V2 interactions on timescales within a trial, we subtracted the mean across time bins within each trial from each neuron. This step removed activity that fluctuated on slow timescales from one stimulus presentation to the next. We then applied DLAG to each dataset separately.

### Intra-areal and subsampled population comparisons

To contrast with the V1-V2 results, we also used DLAG to characterize the interactions between two V1 subpopulations. For each dataset, we randomly split V1 into two equally sized subpopulations (for datasets with an odd number of V1 neurons, we discarded one neuron at random). Each subpopulation was labeled arbitrarily as either “V1a” or “V1b” (Fig. 4c). We then applied DLAG to dissect these V1a-V1b interactions in a manner identical to V1-V2 (Fig. 5, Fig. 6).

We also sought to understand the extent to which the V1-V2 results were driven by disparities in population size between V1 and V2 (Supplementary Fig. 10). For each dataset, we therefore randomly subsampled the V1 population to match the size of the V2 population. We then applied DLAG to each subsampled dataset in the same manner as above.

### Variance explained by DLAG latent variables

After fitting a DLAG model to each experimental dataset, we sought to compare the relative strengths of across- or within-area latent variables extracted from the same dataset (as in Fig. 5) and across different datasets (as in Fig. 6b). To quantify these comparisons, we computed the variance each latent variable explained, as derived from fitted model parameters. From equation (1), the total variance in area i simplifies to

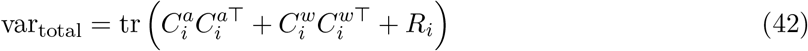

By inspection, the total variance decomposes into three separable components: 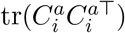, the variance due to across-area activity; 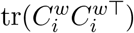, the variance due to within-area activity; and tr(*R_i_*), the variance that is independent to each neuron. In fact, the across-area and within-area components can be decomposed further into contributions by individual latent variables. Let 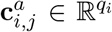 be the *j*^th^ column of 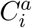, and 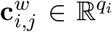 be the *j*^th^ column of 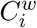. Then, 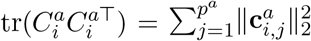, and 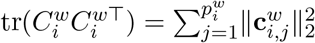.

Because we were interested in variance shared among neurons, rather than independent to each neuron, we focused on the variance components involving 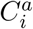 and 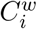, rather than *R_i_*. Furthermore, since the total variance of recorded neural activity may vary widely across animals, stimuli, and recording sessions, we computed two normalized metrics to facilitate comparison of these shared variance components across datasets. First, let c_*i*,*j*_ be the *j*^th^ column of *C_i_*, where 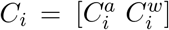 is the same as in equation (12). To visualize the relative strength of latent variables in each area (Fig. 5), we computed

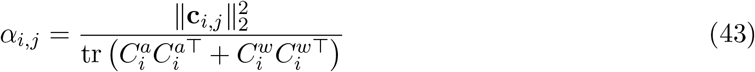

that is, the fraction of shared variance explained by latent variable *j* in area *i*. We then displayed latent time courses multiplied by the appropriate *α*_*i*,*j*_ at each time point. Similarly, to quantify the strength of across-area activity (relative to within-area activity) in each area (Fig. 6b), we computed

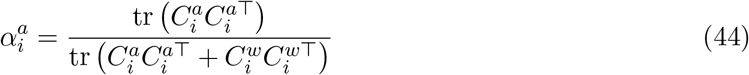

that is, the fraction of shared variance explained by all across-area latent variables in area *i*.

### Uncertainty of estimated delays

DLAG’s performance on the synthetic data presented here suggests that time delays are estimated with high accuracy and precision. For our neural recordings, however, where no “ground truth” is accessible, we sought to assess the certainty with which fitted delay parameters were indeed positive or negative—indicating a particular direction of inter-areal signal flow. We therefore developed the following nonparametric bootstrap procedure.

First, consider a DLAG model that has been fit to a particular dataset with *N* trials. We construct a bootstrap sample *b* = 1,…, *B* from this dataset by selecting *N* trials uniformly at random with replacement (here we used *B* = 1,000). Then, let *ℓ_b_* be the data log-likelihood of the DLAG model evaluated on bootstrap sample *b*. And let *ℓ*_*b*,*j*=0_ be the data log-likelihood of the same DLAG model evaluated on bootstrap sample *b*, but for which *D_j_*, the delay for across-area latent variable *j*, has been set to zero (all other model parameters remain unaltered).

To compare the performance of this “zero-delay” model to the performance of the original model, we define the following statistic:

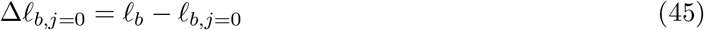

If the zero-delay model performed at least as well as the original DLAG model (equivalently, Δ*ℓ*_*b*,*j*=0_ ≤ 0) on 5% or more of the bootstrap samples, then we could not say, with sufficient certainty, that the delay for across-area variable *j* was strictly positive or strictly negative. Otherwise, we took the magnitude of the delay for across-area variable *j* to differ significantly from zero.

For each of our V1-V2 datasets, then, this procedure allowed us to label some delays as “ambiguous,” where the corresponding population signal could not be confidently categorized as flowing in one direction or the other (Fig. 6c). Finally, note that the concept of ambiguity defined here is distinct from the concept of a variable’s importance in describing observed neural activity: for example, an across-area variable with an ambiguous time delay between areas could, in principle, still explain a large portion of an area’s shared variance.

## Data availability

V1-V2 data are available at the CRCNS data sharing website, at https://doi.org/10.6080/K0B27SHN.

## Code availability

All methods and data analyses described here were implemented and carried out in Matlab (The Mathworks, Inc.). Implementations of DLAG in Matlab and Python will be made publicly available upon publication.

## Acknowledgements

This work was supported by the Dowd Fellowship (E.G.), Simons Collaboration on the Global Brain 542999 (A.K.), 543009 (C.K.M.), 543065 (B.M.Y.), 364994 (A.K., B. M.Y.), NIH R01 EY028626 (A.K.), NIH U01 NS094288 (C.K.M.), NIH R01 HD071686 (B.M.Y.), NIH CRCNS R01 NS105318 (B.M.Y.), NSF NCS BCS 1533672 and 1734916 (B.M.Y.), NIH CR-CNS R01 MH118929 (B.M.Y.), and NIH R01 EB026953 (B.M.Y.).

## Author contributions

E.G., A.I.J., J.D.S., A.K., C.K.M., and B.M.Y. designed the analyses. E.G. implemented code and performed all analyses. A.Z. and A.K. designed and performed the experiments. E.G., A.K., C.K.M., and B.M.Y. wrote the manuscript. E.G., A.I.J., J.D.S., A.K., C. K.M., and B.M.Y. edited the manuscript. A.K., C.K.M., and B.M.Y. contributed equally to this work.

## Supplementary information

**Supplementary Figure 1.**
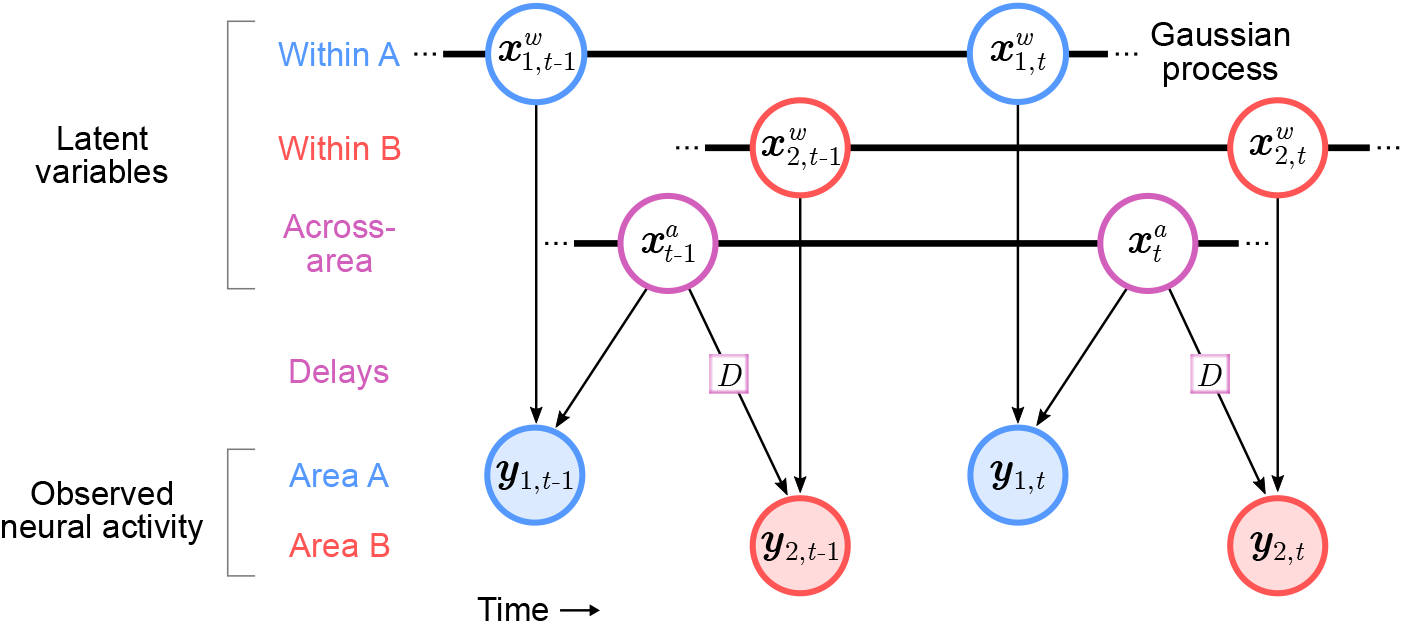
DLAG directed graphical model representation. Filled circles represent observed variables (i.e., observed neural activity in each area), where y_1,*t*_ and y_2,*t*_ are the observed neural activity in area A and B, respectively, at time *t*. Unfilled circles represent latent variables, where 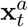 are across-area variables at time *t*; 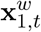 and 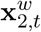 are within-area variables in area A and B, respectively, at time *t*. *D* represents the set of relative time delay parameters between the two areas. Color indicates a variable’s or parameter’s association with area A (blue), area B (red), or both (magenta). Arrows indicate conditional dependence relationships between variables. In particular, the arrows point from latent variables to observed neural activity, framing DLAG as a generative model. Thick black lines indicate that variables are related in time via a Gaussian process. Here two time steps are shown (*t* – 1 and *t*), and time evolves from left to right.

**Supplementary Figure 2.**
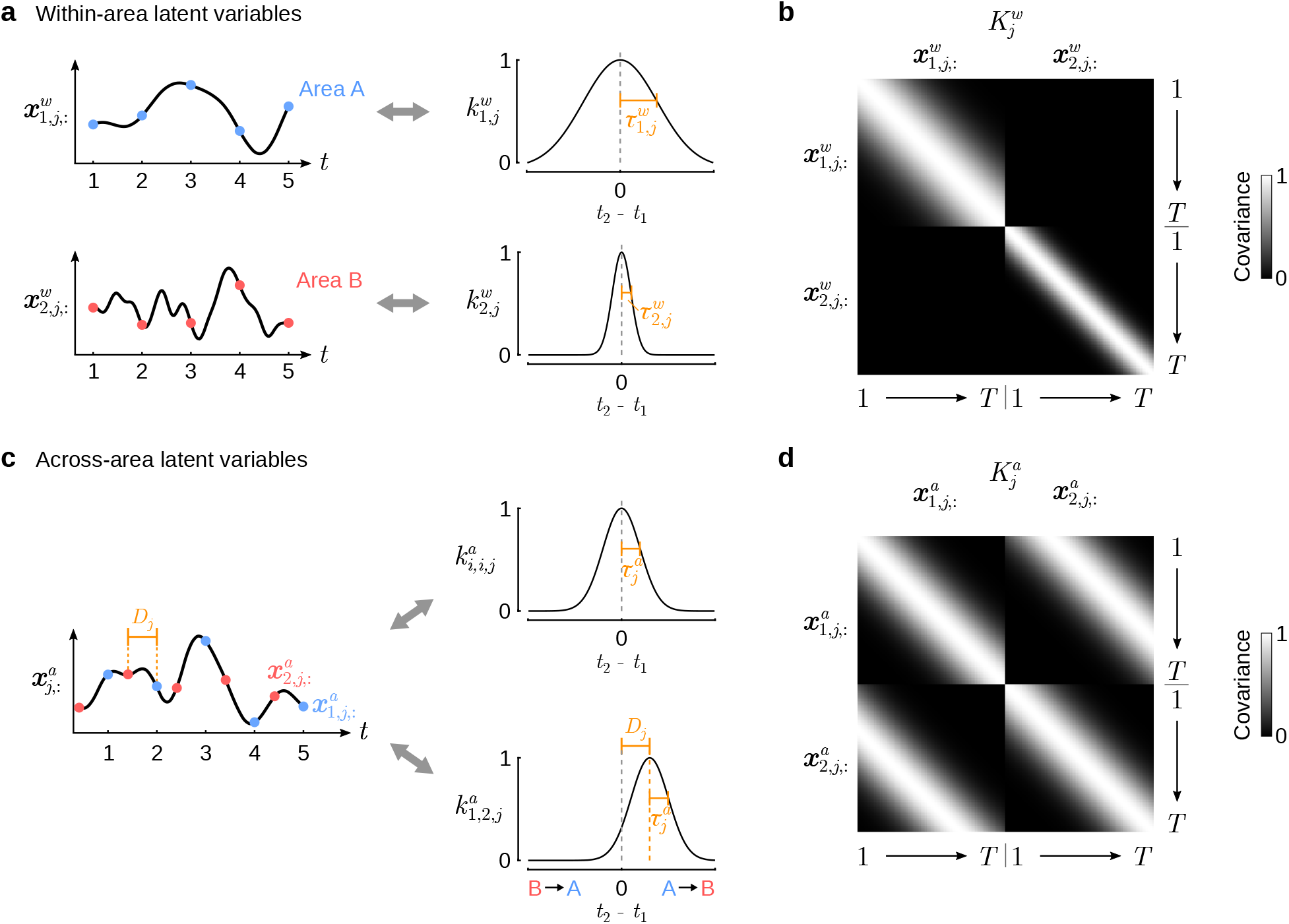
Use of Gaussian processes in the DLAG state model. (**a**) Within-area state model. Left column: Within-area time courses (area A: 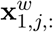, blue points; area B: 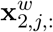, red points) can be described as a finite number of samples drawn from a Gaussian process (GP) for each area and each *j*. Right column: The temporal structure of each within-area GP is governed by a covariance function (area A: 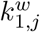; area B: 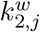). The squared exponential (SE) function, chosen for the present work, is defined by a timescale parameter 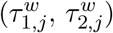, which controls the width the covariance kernel, or equivalently, how quickly the latent variable changes over time. (**b**) An example set of within-area GP covariance matrices 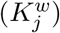. The banded structure emerges from the choice of squared exponential function and stationarity of the GP covariance. Note the independence of within-area latent variables across areas: each latent variable has its own characteristic timescale, and cross-covariance terms are all zero. (**c**) Across-area state model. Left column: Like within-area time courses, across-area time courses can also be described as a finite number of samples drawn from a GP. In contrast to the within-area time courses, which are independent across areas, across-area time courses are coupled across areas, drawn from a common GP 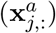. The sampling grid of area A (blue) is shifted by a time delay (*D_j_*) relative to that of area B (red). Right column: The temporal structure of the common GP is governed by a SE covariance function. The width of the auto- and cross-covariances (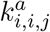 and 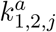, respectively) is controlled by a timescale parameter 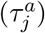. The center of the cross-covariance is controlled by the delay parameter *D_j_* (positive delays: A leads B; negative delays: B leads A). (**d**) An example across-area GP covariance matrix 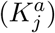. The banded structure emerges from the choice of squared exponential function and stationarity of the GP covariance. Note the non-zero cross-covariance terms in the off-diagonal blocks of 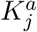: the banded structure is shifted from the diagonal of each off-diagonal block by the delay parameter *D_j_*.

**Supplementary Figure 3.**
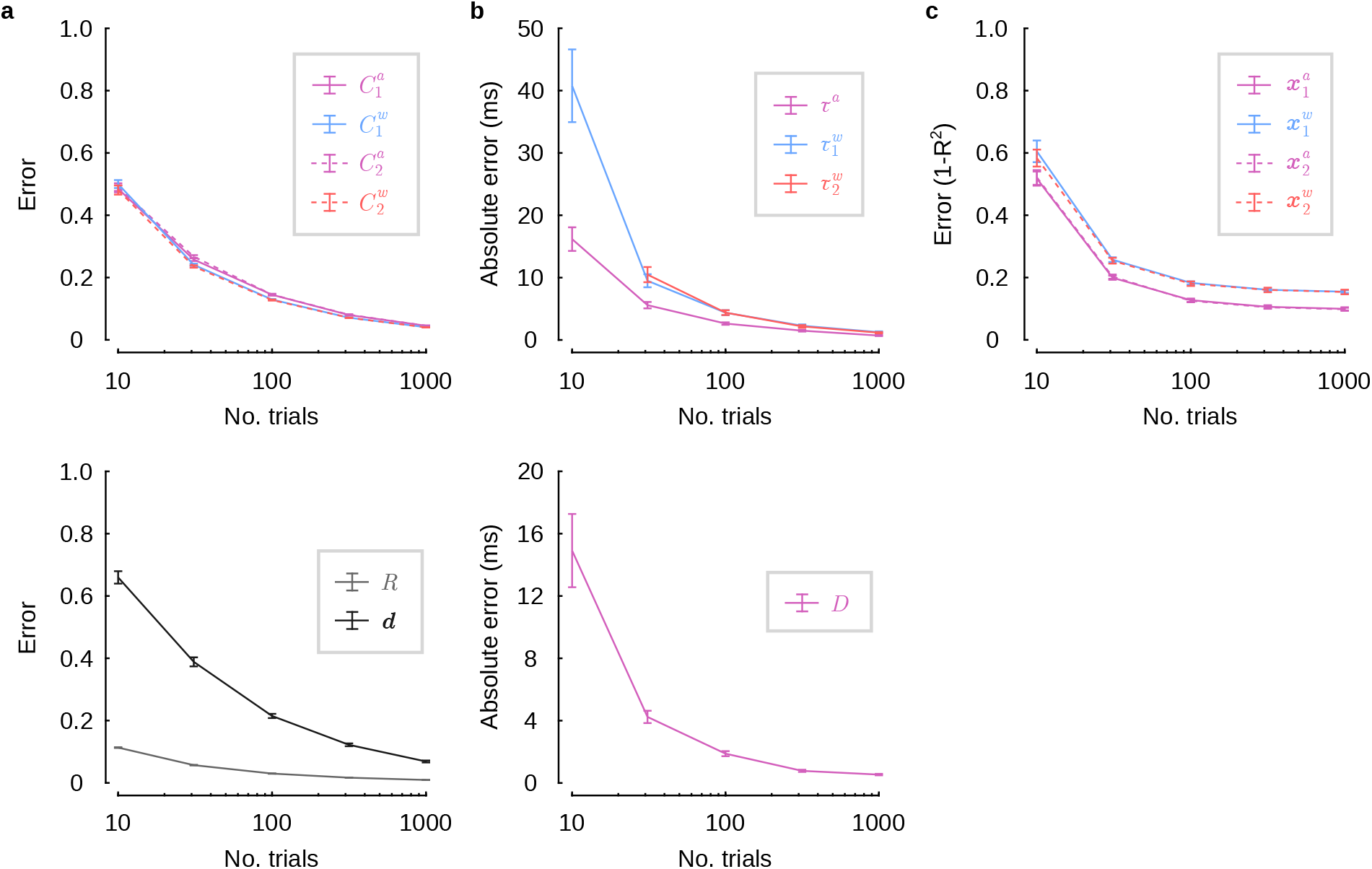
DLAG performance improves with increasing number of trials. We sought to characterize DLAG’s performance as a function of the number of available trials. We therefore synthesized 25 additional datasets (via the DLAG generative model) with the following characteristics: *N* = 1000 trials; *q*_1_ = *q*_2_ = 50 neurons per area; 500 ms trial lengths with 20 ms sampling period (for *T* = 25 samples per trial); latent dimensionalities 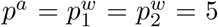; signal-to-noise ratios 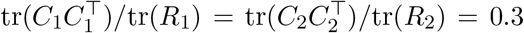; GP timescales *τ^a^*, 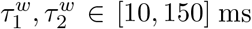; and delays *D* ∈ [-30,30] ms. We took subsets of trials from these datasets, and fit DLAG to increasingly large subsets (sizes equally spaced on a log scale from 10 to 1000 trials). (**a**) Error of observation model parameter estimates versus number of trials. Top: Error of across- and within-area loading matrix estimates decreases with increasing number of trials. (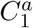: solid magenta; 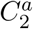: dashed magenta; 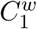: solid blue; 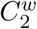: dashed red). Bottom: Error of covariance matrix (*R*: light gray) and mean parameter (**d**: dark gray) estimates decreases with increasing number of trials. Error bars represent SEM across 25 independent simulated datasets. (**b**) Absolute error (in ms) of state model parameter estimates versus number of trials. Top: The error in across- and within-area GP timescale estimates decreases as number of trials increases (*τ^a^*: magenta; 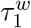: blue; 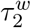: red). Error of within-area timescale estimates 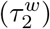 have been omitted for values of 10 trials, where absolute error was 212.1±174.4 ms (mean and SEM across all within-area timescales). Given insufficient statistical power, some GP timescale estimates (likely for latent dimensions that explain little shared variance within an area) become large (i.e., larger than the length of a trial)—to the point where smoothed population activity in the corresponding dimension is effectively constant within a trial. Bottom: The error in delay parameter estimates decreases as number of trials increases. Error bars represent SEM across 125 latent variables. (**c**) Error (1 – *R*^2^) of latent variable time course estimates versus number of trials. Moment-by-moment estimates of latent variable time courses improve as the number of trials increases (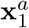: solid magenta; 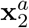: dashed magenta; 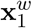: solid blue; 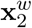: dashed red). Error bars represent SEM across 25 independent simulated datasets.

**Supplementary Figure 4.**
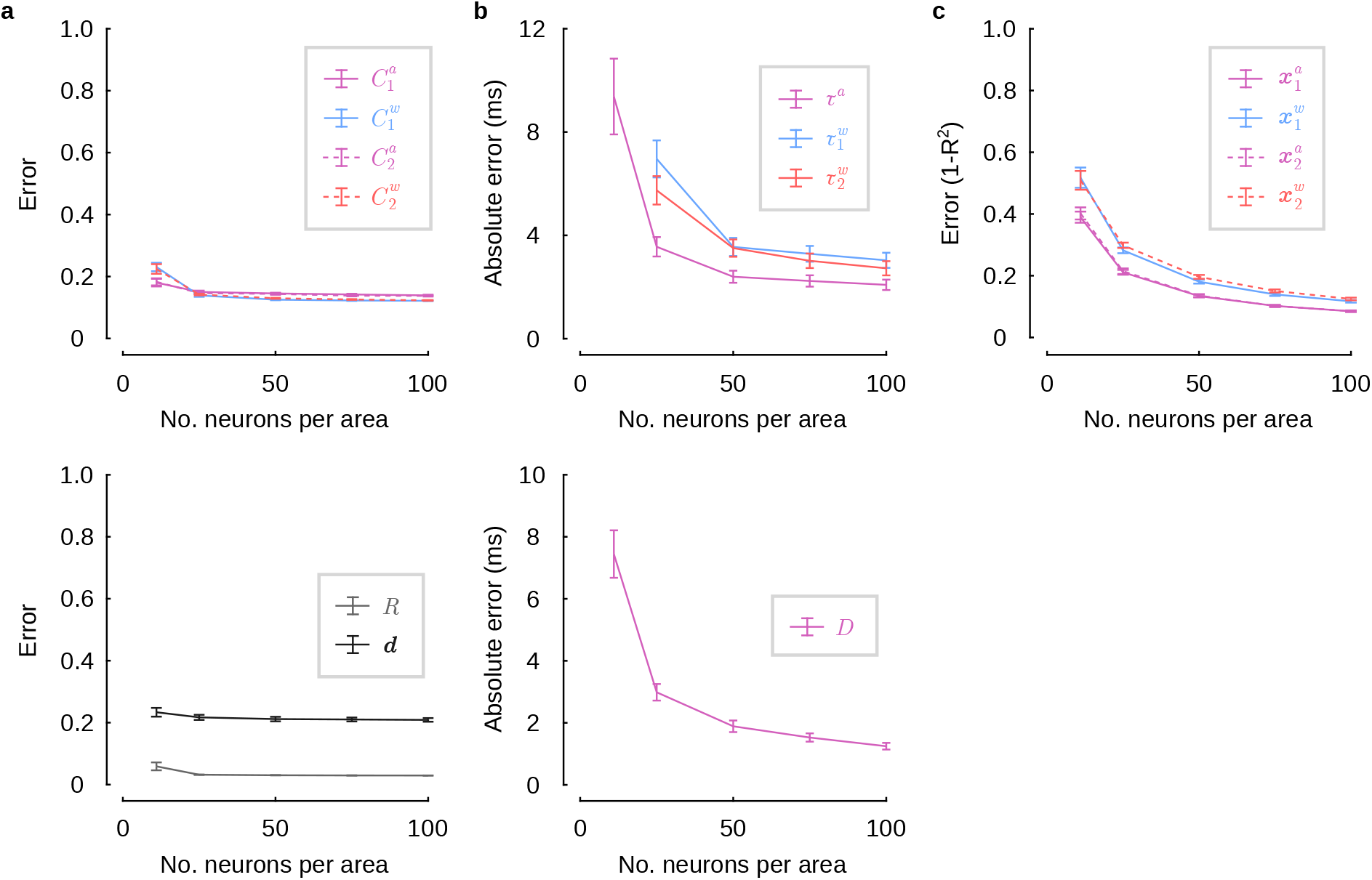
DLAG performance improves with increasing number of neurons (and fixed latent dimensionality). We sought to characterize DLAG’s performance as a function of the number of recorded neurons. We therefore synthesized 25 additional datasets (via the DLAG generative model) with the following characteristics: *N* = 100 trials; *q*_1_ = *q*_2_ = 100 neurons per area; 500 ms trial lengths with 20 ms sampling period (for *T* = 25 samples per trial); latent dimensionalities 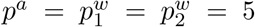; signal-to-noise ratios 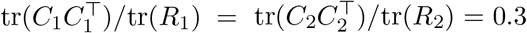; GP timescales *τ^a^*, 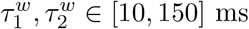; and delays *D* ∈ [-30, 30] ms. We took subsets of neurons from these datasets, and fit DLAG to increasingly large subsets (11, 25, 50, 75, and 100 neurons in each area). (**a**) Error of observation model parameter estimates versus number of neurons per area. Top: Error of across- and within-area loading matrix estimates decreases with increasing number of neurons per area. (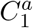: solid magenta; 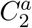: dashed magenta; 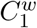: solid blue; 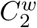: dashed red). Bottom: Error of covariance matrix (*R*: light gray) and mean parameter (**d**: dark gray) estimates decreases with increasing number of neurons per area. Error bars represent SEM across 25 independent simulated datasets. (**b**) Absolute error (in ms) of state model parameter estimates versus number of neurons per area. Top: The error in across- and within-area GP timescale estimates decreases as number of neurons per area increases (*τ^a^*: magenta; 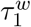: blue; 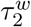: red). Error of within-area timescale estimates have been omitted for values of 11 neurons per area, where absolute error was 60.1±39.2 ms for 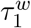 and 93.7±46.3 ms for 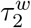 (mean and SEM across all within-area timescales). Given insufficient statistical power, some GP timescale estimates (likely for latent dimensions that explain little shared variance within an area) become large (i.e., larger than the length of a trial)—to the point where smoothed population activity in the corresponding dimension is effectively constant within a trial. Bottom: The error in delay parameter estimates decreases as number of neurons per area increases. Error bars represent SEM across 125 latent variables. (**c**) Error (1 – *R*^2^) of latent variable time course estimates versus number of neurons per area. Moment-by-moment estimates of latent variable time courses improve as the number of neurons per area increases (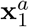: solid magenta; 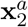: dashed magenta; 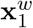: solid blue; 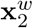: dashed red). Error bars represent SEM across 25 independent simulated datasets.

**Supplementary Figure 5.**
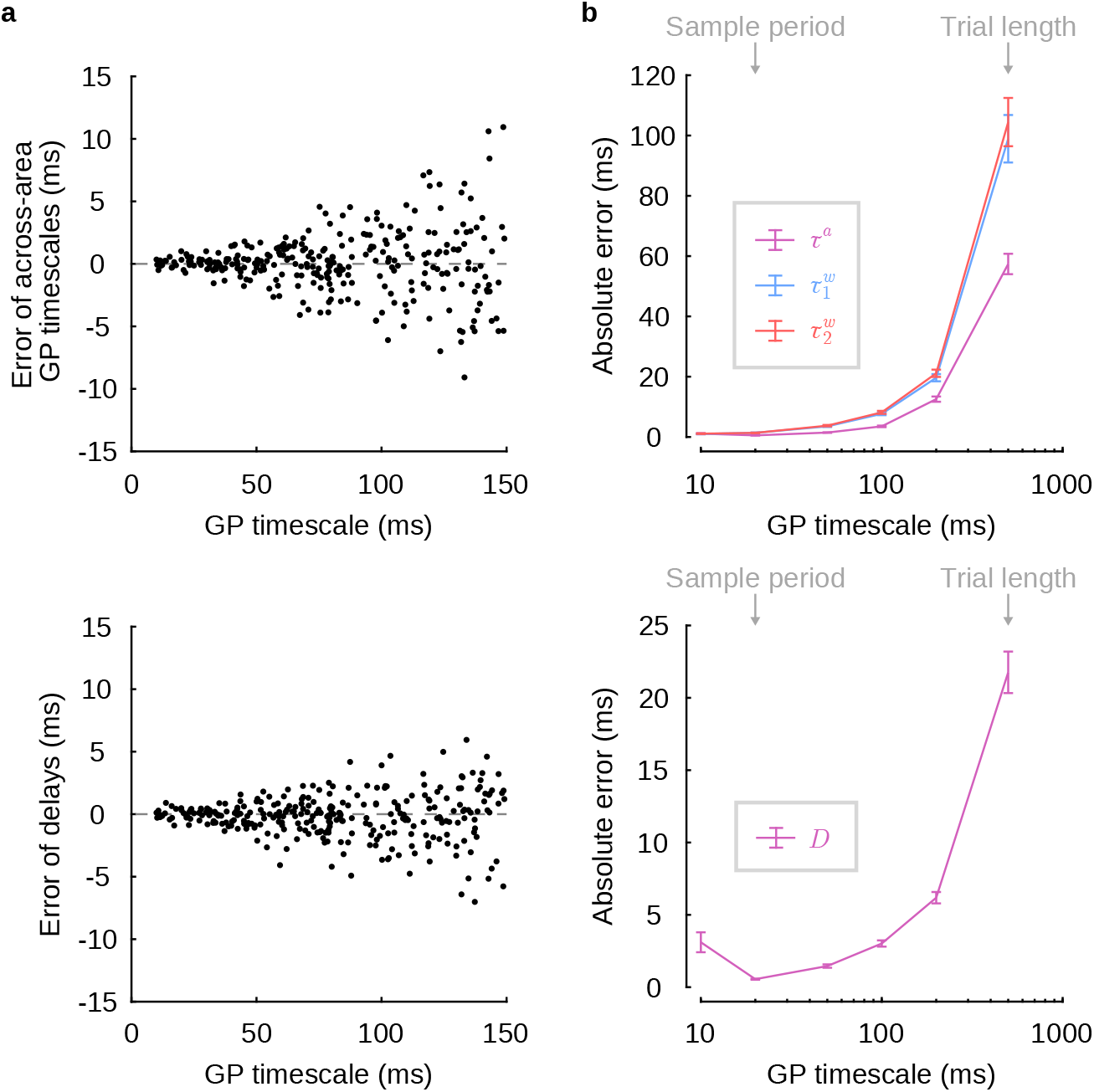
Uncertainty of DLAG timescale and delay estimates increases with increasing latent timescale. (**a**) Error (in ms; estimate minus ground truth value) of across-area GP timescale (top) and delay (bottom) estimates for each latent variable shown in Fig. 3c,d. The variance of both GP timescale and delay estimates appears to increase as the underlying ground truth GP timescale increases. For intuition, consider the extreme case of a latent variable whose time course is constant, or equivalently, whose autocovariance function (Supplementary Fig. 2) is flat. Then, a range of DLAG models with any delay and any sufficiently long GP timescale could explain the data equally well. Similarly, as the GP timescale of a latent variable increases, a wider of range of delay and timescale estimates leads to similar model performance, particularly in the presence of noise. (**b**) To verify the apparent trend in (a), we systematically characterized the accuracy of GP timescale and delay parameter estimates as a function of ground truth GP timescale. We synthesized additional datasets (via the DLAG generative model) with the following characteristics: *N* = 100 trials; *q*_1_ = *q*_2_ = 50 neurons per area; 500 ms trial lengths with 20 ms sampling period (for *T* = 25 samples per trial); latent dimensionalities 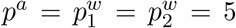; signal-to-noise ratios 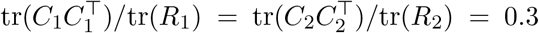; and delays *D* ∈ [-30, 30] ms. Each dataset’s within- and across-area latent variables were given the same GP timescale; and across 150 datasets, we considered six different timescales (25 datasets synthesized for each timescale), ranging in length from half the sampling period to the length of the trial (10 ms, 20 ms, 50 ms, 100 ms, 200 ms, 500 ms). Top: Absolute error (in ms) of across- and within-area GP timescale estimates increases as underlying GP timescale increases (*τ^a^*: magenta; 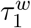: blue; 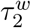: red). Bottom: Absolute error (in ms) of delay parameter estimates increases as underlying GP timescale increases. The lowest error is achieved when GP timescales are equal to the sampling period of observations. For GP timescales larger than the sampling period, the error increases according to the intuition outlined above. For GP timescales less than the sampling period, error increases because latent variable time courses change faster than can be accurately observed at that sampling rate, thereby introducing uncertainty in estimates. Error bars represent SEM across 125 latent variables.

**Supplementary Figure 6.**
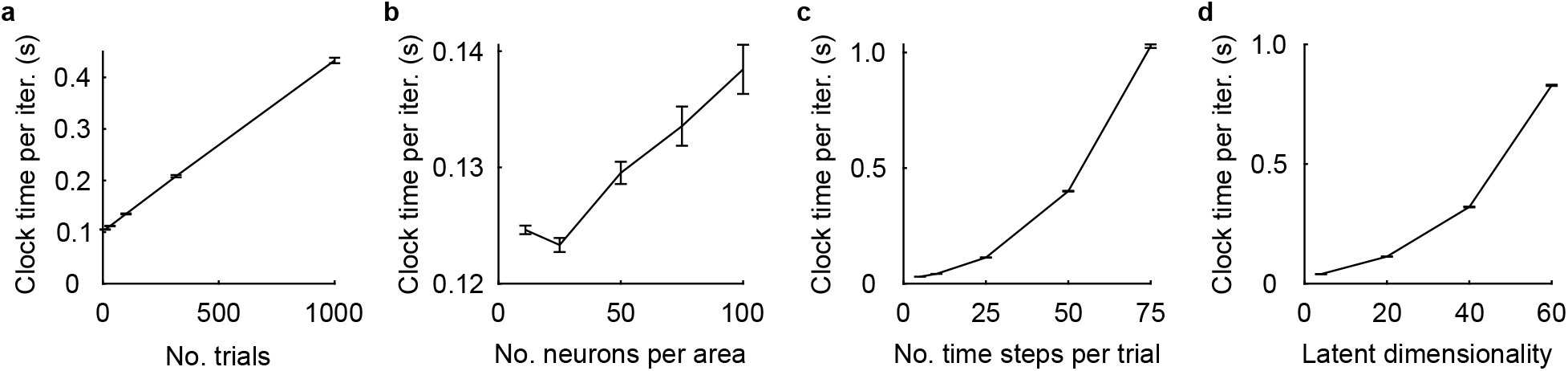
DLAG runtime as a function of number of trials, number of neurons, trial length, and latent dimensionality. (**a**) The average clock time (in seconds) per DLAG EM iteration scales (approximately) linearly with the number of trials. These runtime analyses were carried out on synthetic datasets with *q*_1_ = *q*_2_ = 50 neurons in each area; *T* = 25 time steps per trial; and latent dimensionalities 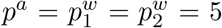 (total number of latent dimensions 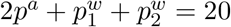). (**b**) The average clock time (in seconds) per DLAG EM iteration scales (approximately) linearly with the number of neurons per area. These runtime analyses were carried out on synthetic datasets with *N* = 100 trials; *T* = 25 time steps per trial; and latent dimensionalities 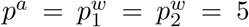 (total number of latent dimensions 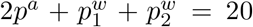). (**c**) The average clock time (in seconds) per DLAG EM iteration scales (approximately) quadratically with the number of time steps per trial. Runtime scales quadratically, rather than linearly (as in (a)), because DLAG describes the temporal structure within each trial via Gaussian processes. These runtime analyses were carried out on synthetic datasets with *N* = 100 trials; *q*_1_ = *q*_2_ = 50 neurons in each area; and latent dimensionalities 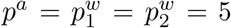 (total number of latent dimensions 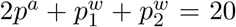). (**d**) The average clock time (in seconds) per DLAG EM iteration scales (approximately) quadratically with the total number of latent dimensions 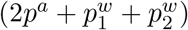. These runtime analyses were carried out on synthetic datasets with *N* = 100 trials; *q*_1_ = *q*_2_ = 50 neurons in each area; and *T* = 25 time steps per trial. In (a)-(d), error bars represent SEM across 25 independent simulated datasets. Results were obtained on a Red Hat Enterprise Linux machine (release 7.9, 64-bit) with 250GB of RAM running Matlab (R2019a), on an Intel Xeon CPU (E5-2695 v3, 2.3 GHz).

**Supplementary Figure 7.**
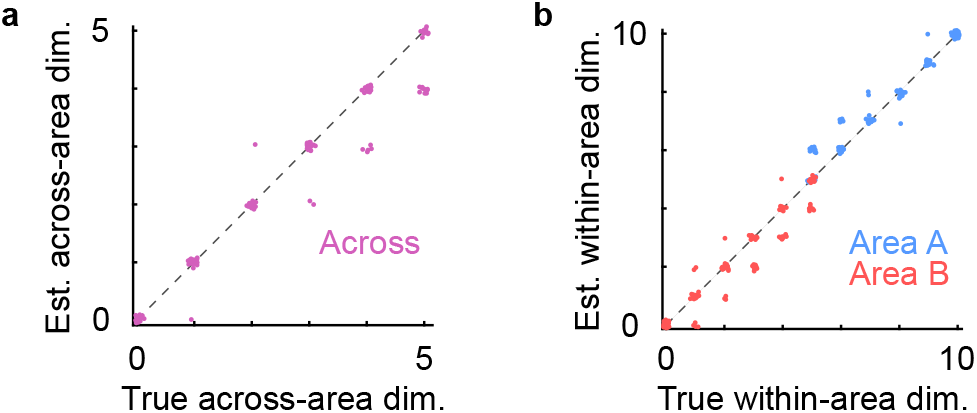
Model selection remained accurate in a regime of weak shared variance. To explore a scenario in which our model selection procedure began to misestimate within- or across-area dimensionalities, we synthesized an additional 120 independent datasets in which the strength of activity due to latent variables (i.e., shared variance in each area) relative to the strength of variance independent to each neuron (i.e., signal-to-noise ratio) was smaller than we observed in the V1-V2 recordings. Specifically, we generated datasets from the DLAG generative model such that the ratio 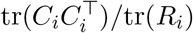 was 0.1 for each area *i* (compared to 0.3 and 0.2 in area A and area B, respectively, in the original synthetic datasets; see Methods). All other data characteristics remained the same as in the original data. (**a**) Estimated versus ground truth across-area dimensionalities. (**b**) Estimated versus ground truth within-area dimensionalities. For both (a) and (b), data points are integer-valued, but randomly jittered to show points that overlap. Estimated across- and within-area dimensionalities never deviated from the ground truth by more than one, and the datasets with an absence of across- or within-area structure were always correctly identified. On datasets where a mismatch between estimated and ground truth dimensionalities did occur, the primary source of that inaccuracy originated from the initial factor analysis (FA) stage of model selection, rather than the second stage involving DLAG. Of the 66 datasets in which FA correctly estimated the total dimensionalities in both area A and area B, 65 datasets were correctly estimated following the second stage of model selection with DLAG models; and, irrespective of the accuracy after the FA stage, across-area dimensionality was correct on 101 of 120 datasets. This phenomenon might be expected, given the mismatch between the FA model and the DLAG generative model used to synthesize these datasets. It also provides the opportunity to explore the impact of imperfect dimensionality estimates—inevitable in real data—on the estimation and interpretation of DLAG’s parameters and latent variables following fitting (see Supplementary Fig. 8, Supplementary Fig. 9).

**Supplementary Figure 8.**
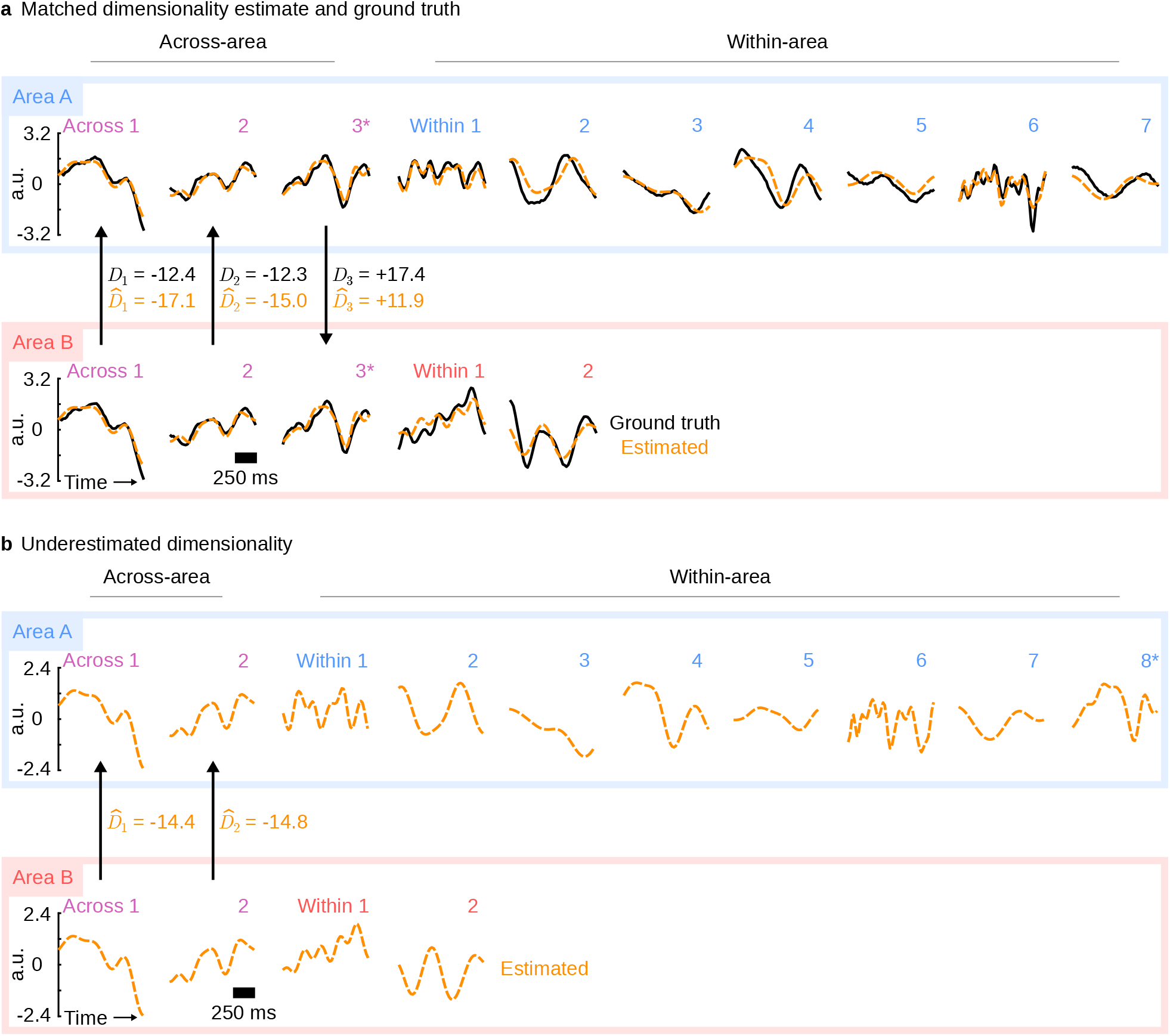
DLAG’s parameter and latent variable estimates remained stable when dimensionality was underestimated. Here we present a case study from one of the synthetic datasets presented in Supplementary Fig. 7, in which the total dimensionality of area B was underestimated during the initial factor analysis (FA) model selection stage, and across-area dimensionality was underestimated in the second stage. (**a**) For reference, we first fit a DLAG model with the correct number of within- and across-area latent variables, i.e, no model selection was performed. With real data, we would not have access to this information, but here we use it to understand the real data scenario in (b). Shown are single-trial latent-variable time course estimates produced by the fitted model along with the ground truth (one example trial shown). Top row / blue box: area A; bottom row / red box: area B. Left: across-area; right: within-area. Orange dashed traces: DLAG estimates; black solid traces: ground truth. a.u.: arbitrary units. Delays reported in ms. Even in the weak-shared variance regime, estimates are qualitatively close to the ground truth. The asterisks (‘*’) are intended to highlight the third across-area latent variable for each area, which becomes mistaken as a within-area A latent variable when area B’s dimensionality is underestimated (see within-area A latent variable 8 in (b)). (**b**) We next consider the model chosen through model selection, as we would with real data. The estimated number of latent variables in area B and the estimated number of across-area variables were each one fewer than the respective ground truth. Shown are single-trial latent-variable time course estimates produced by this model (same trial shown as in (a)). Qualitatively, time course estimates closely match those of the model in (a), in which the correct number of within- and across-area variables was used (compare estimated latent variables with the same index across (a) and (b)). Furthermore, delay estimates are only slightly affected. By inspection, the third across-area latent variable pair (marked by the asterisks in (a)) now appears as the eighth within-area A latent variable (also marked by an asterisk). Note that the ordering of latent variables is arbitrary; we have ordered the latent variables here to facilitate visual illustration.

**Supplementary Figure 9.**
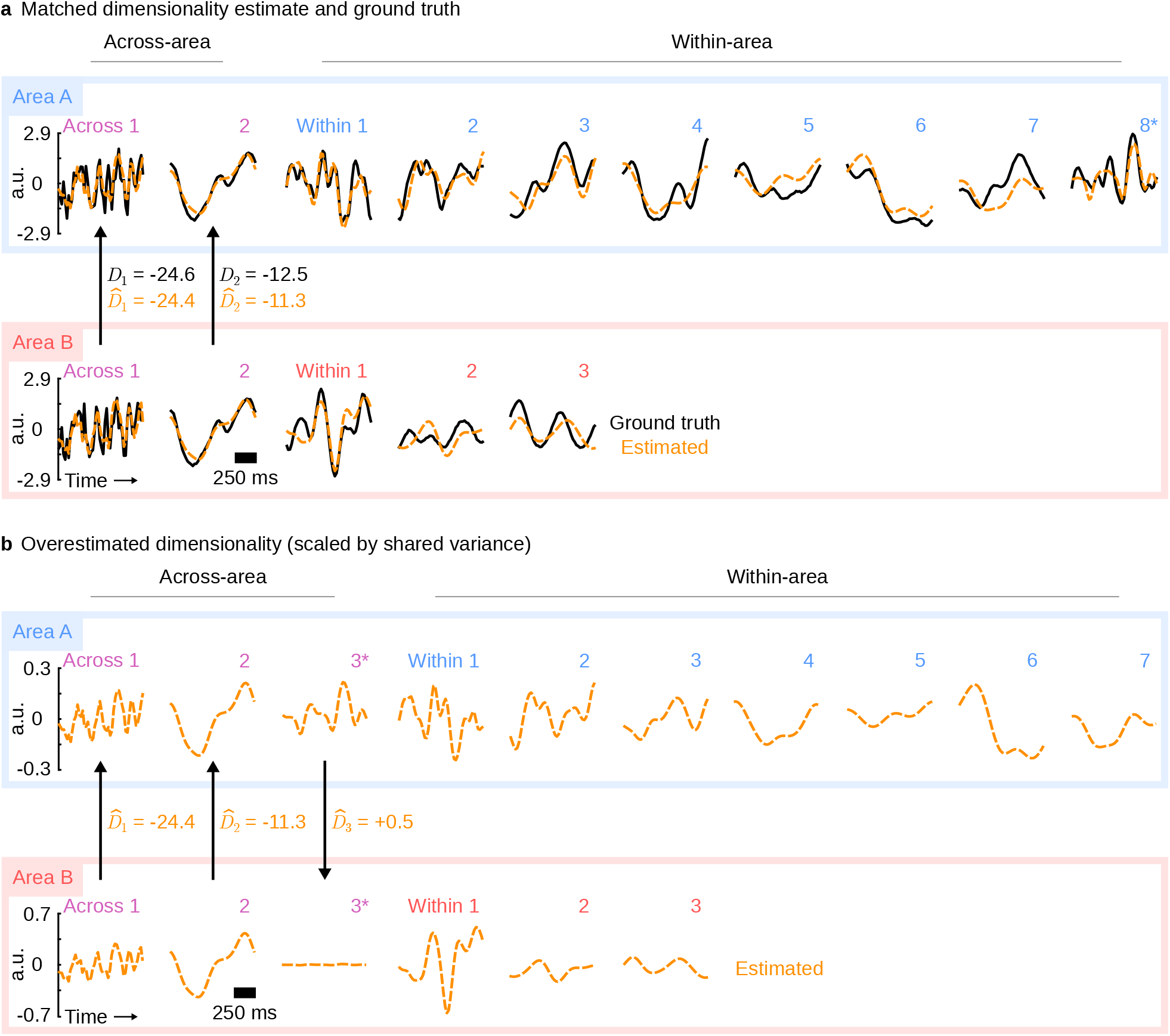
DLAG’s parameter and latent variable estimates remained stable when dimensionality was overestimated. Here we present a case study from one of the synthetic datasets presented in Supplementary Fig. 7, in which the total dimensionality of area B was over-estimated during the initial factor analysis (FA) model selection stage, and across-area dimensionality was overestimated in the second stage. (**a**) For reference, we first fit a DLAG model with the correct number of within- and across-area latent variables, i.e., no model selection was performed. With real data, we would not have access to this information, but here we use it to understand the real data scenario in (b). Shown are single-trial latent-variable time course estimates produced by the fitted model along with the ground truth. Same conventions as in Supplementary Fig. 8. Even in the weak-shared variance regime, estimates are qualitatively close to the ground truth. The asterisk (‘*’) is intended to highlight the eighth within-area A latent variable, which becomes mistaken as an across-area variable when area B’s dimensionality is overestimated (see across-area variable 3 in (b)). (**b**) We next consider the model chosen through model selection, as we would with real data. The estimated number of latent variables in area B and the estimated number of across-area variables were each one more than the respective ground truth. Shown are single-trial latent-variable time course estimates produced by this model (same trial shown as in (a)). Qualitatively, time course estimates closely match those of the model in (a), in which the correct number of within- and across-area variables was used (compare estimated latent variables with the same index across (a) and (b)). By inspection, the eighth within-area A latent variable (marked by the asterisk in (a)) now appears as the third across-area latent variable (also marked by asterisks). This phenomenon is straightforward to diagnose: here, we have additionally scaled each latent variable by the fraction of shared variance it explains within its respective area (see Methods; same convention as in Fig. 5). The third across-area latent variable explains little shared variance in area B, consistent with the ground truth. Note that the ordering of latent variables is arbitrary; we have ordered the latent variables here to facilitate visual illustration.

**Supplementary Figure 10.**
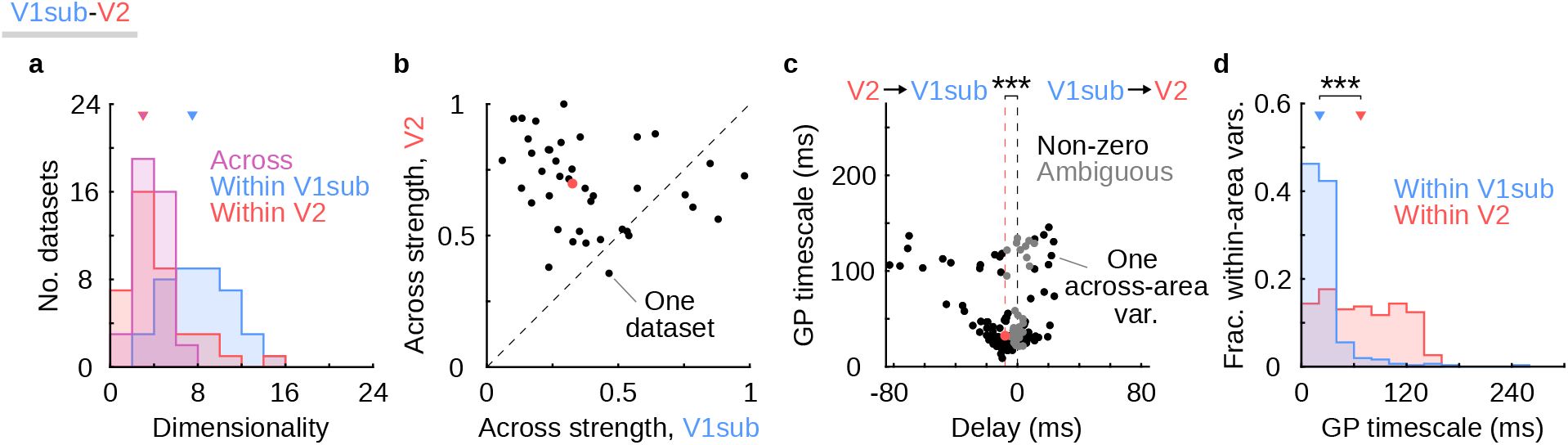
V1-V2 results are preserved when V1 is subsampled to match V2 in population size. Same conventions as in Fig. 6. We sought to understand the extent to which the results reported in Fig. 6 were driven by the fact that V1 populations were larger than V2 populations. All else being equal, more neurons allows one to reliably identify more latent dimensions [70]. For each dataset, we thus randomly subsampled the V1 population (‘V1sub’) to match the size of the V2 population. We then applied DLAG to each subsampled dataset in the same manner as in Fig. 6. (**a**) V1sub-V2 within- and across-area dimensionalities. Compared to Fig. 6, median across-area dimensionality (3) was the same. As a consequence of the smaller population size, median within-V1sub dimensionality (7.5) decreased, but remained higher than median across-area and median within-V2 (3) dimensionalities. Within-V2 dimensionality was 0 in 1 of 40 datasets. (**b**) Fraction of shared variance of each area explained by across-area latent variables in V1sub and in V2. Despite population sizes now being the same, across-area strength is still significantly greater in V2 than in V1sub (median V1sub: 0.33; median V2: 0.70; onesided paired sign test; *p* < 0.001), as in Fig. 6. Even after controlling for V1 population size, the within-area dimensionality of V1sub and V2 are not equal. It is possible that the difference in across-area strength seen in (b) is implied by, and therefore redundant with, the difference in within-area dimensionalities seen in (a). Specifically, the weaker across-area strength in V1 relative to V2 might be implied by the greater number of within-V1sub dimensions relative to the number of across-area dimensions. To test this possibility, we recomputed the median across-area strengths for V1 and V2, considering only datasets such that the distributions of within-V1sub and within-V2 dimensionalities were the same. Sixteen datasets remained after this distribution-matching procedure (the 16 datasets for V1 were not necessarily the same 16 datasets as for V2). The medians in V1 and V2 were nearly unchanged (V1sub: 0.33; V2: 0.67). Across-area strengths therefore convey a difference in the properties of V1 versus V2 activity that could not be seen from differences in dimensionality alone. (c) Gaussian process (GP) timescale vs. time delay for across-area latent variables. Across all 40 datasets, the delays of 97 of 136 across-area variables were deemed significantly non-zero, and the remaining 39 delays were deemed ambiguous. These values are nearly identical to those reported in Fig. 6. Similarly, delays remained significantly less than zero, representing feedback interactions from V2 to V1sub (median delay across all significantly non-zero across-area variables: −8 ms; ‘***’: one-sided one-sample sign test on ‘non-zero’ delays, *p* < 0.001). Among the significantly non-zero delays, 67% were negative. The magnitude of significant negative delays (median: −12 ms) remained greater than the magnitude of significant positive delays (median: +8ms). (d) GP timescales for within-area latent variables. GP timescales within V1sub and within V2 are similar to those reported in Fig. 6 (median across 307 within-V1sub latent variables: 21 ms; median across 153 within-V2 latent variables: 68 ms). Furthermore, as in Fig. 6, within-V2 GP timescales are significantly longer than within-V1sub GP timescales (‘***’: one-sided Wilcoxon rank sum test, *p* < 0.001).

## Supplementary Discussion

### Statistical tradeoffs between within- and across-area latent variables

Throughout this work, we have described how DLAG decomposes observed neural activity into a linear combination of within- and across-area latent variables. Equivalently, DLAG partitions each area’s population space into distinct within- and across-area subspaces, which represent characteristic ways in which the neurons covary (Fig. 2). Here we investigate more deeply why the within-area latent variables are a necessary model component, even if across-area activity is of primary scientific interest. Toward that end, we will consider an alternative interpretational perspective: namely, that DLAG performs a low-rank decomposition of the covariance matrix of a time series. This alternative perspective also illuminates a general statistical phenomenon—not specific to DLAG—that any multi-area time series method must consider.

### DLAG performs a low-rank covariance decomposition

Let us first express the DLAG model not only for a single time point, as in equation (15), but for all time points in a sequence. In particular, we will collect observed and latent variables in a manner that highlights group structure (i.e., organized differently than in equations (16) and (17)). We define 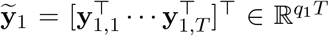 and 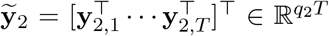, obtained by vertically concatenating the observed neural activity y_1,*t*_ and y_2,*t*_ in areas 1 and 2, respectively, across all times *t* = 1,…, *T*. We collect the across- and within-area latent variables for each area similarly. Let 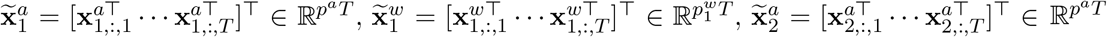, and 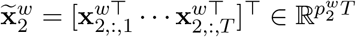.

Then, we rewrite the state and observation models as follows:

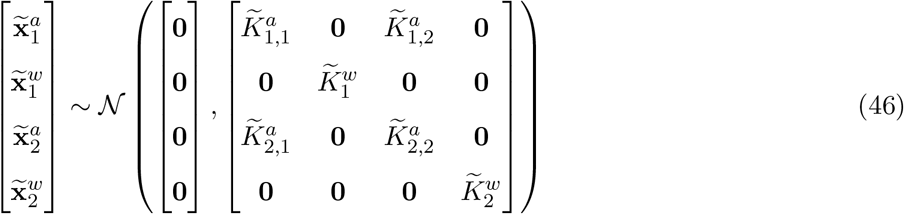

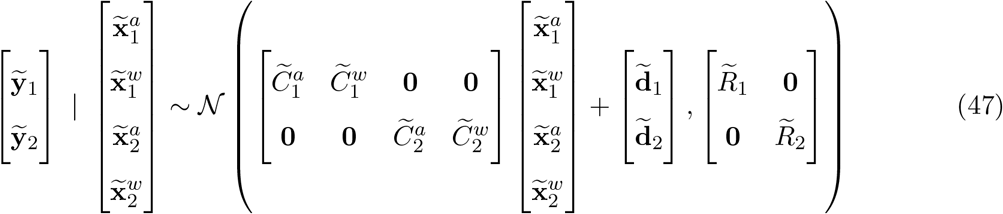

where 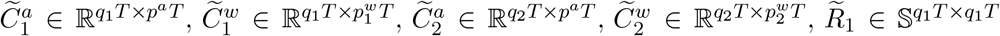, and 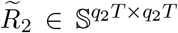 are all block diagonal matrices comprising *T* copies of the loading matrices 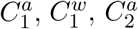, and 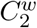, and observation noise covariance matrices *R*_1_ and *R*_2_, respectively. 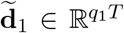 and 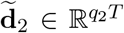 are constructed by vertically concatenating *T* copies of mean parameters d_1_ and d_2_, respectively. Note that equations (46) and (47) above are equivalent to equations (16) and (17), but with variables rearranged.

Each within-area covariance matrix 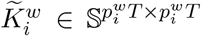, for area *i* = 1, 2 has the following block structure:

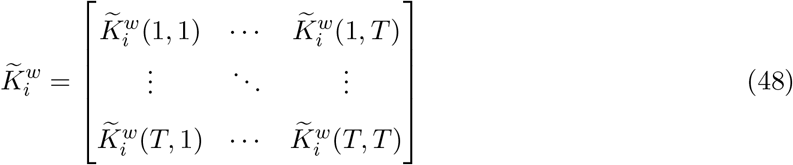

where each block 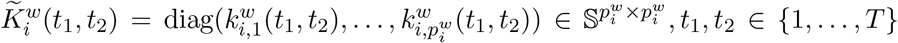 is a diagonal matrix whose elements are computed according to the covariance function defined in equations (4) and (5).

Each across-area auto- or cross-covariance matrix 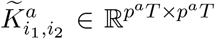, for areas *i*_1_, *i*_2_ ∈ {1, 2} has analogous structure:

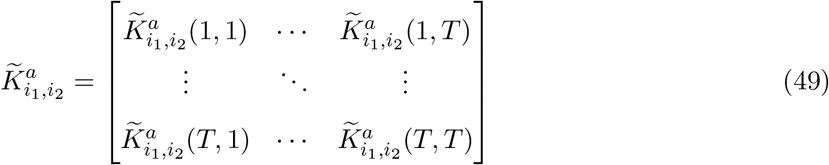

where each block 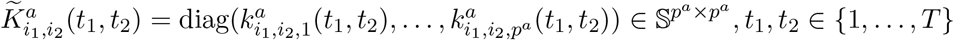 is a diagonal matrix whose elements are computed according to the covariance function defined in equations (7) and (8). Note that the cross-covariance matrices are transposes of one another, i.e., 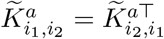.

Upon inspection of equation (46), the statistical dependency between latent variables becomes clear. However, the statistical dependency between observed neural activity in each area, ỹ_1_ and ỹ_2_, is not obvious, since the structure of equation (47) suggests that they might be decoupled. The relationship between observed areas becomes clear when we consider their joint distribution, after marginalizing out the latent variables:

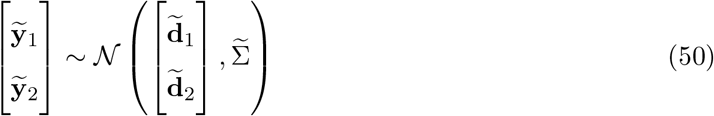

where

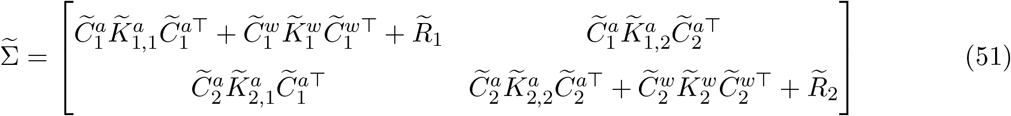

Equation (51) makes explicit the alternative interpretational perspective of DLAG: DLAG performs a low-rank decomposition of the covariance matrix 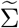. This decomposition is illustrated graphically in Fig. S1a. For simplicity, we illustrate a covariance matrix for areas with three neurons each, over two time points. The shading of blocks of the covariance matrix illustrate which type of DLAG parameter is responsible for explaining that particular portion of covariance (magenta: across-area; blue/red: within-area; gray: independent single-neuron variability). Regions of overlap (i.e., where both blue/magenta or red/magenta shading are present) illustrate portions of covariance that both within- and across-area variables are responsible for explaining. Any regions of white indicate that no model parameters explain that portion of covariance.

The across-area parameters (note the fully magenta-shaded across-area covariance component in Fig. S1a) serve to explain covariance among all neurons, in both areas. Within-area parameters (blue and red shading, for areas 1 and 2, respectively) serve to explain covariance among neurons within each area, but not across areas (note the white across-area blocks for the within-area covariance component). Importantly, the only parameters in the DLAG model capable of explaining covariance across areas are the across-area parameters (only magenta shading is present in the across-area blocks of 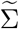). And interestingly, within-area components fully overlap across-area components in the within-area blocks of 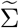, suggesting a potential redundancy. However, as we will discuss below, the overall structure of the decomposition shown in Fig. S1a is critical to the interpretation of across-area variables—that they isolate neural interactions *across* areas (and minimally reflect purely within-area interactions).

### A time series within-area model must accompany a time series across-area model

To build further intuition, let us consider the scenario where within- and across-area covariances are modeled statically—without considering the flow of time (Fig. S1b). Static covariance decompositions result, for example, from the probabilistic canonical correlation analysis (pCCA) model [67], which includes static across-area latent variables and no within-area latent variables (within-area covariance is instead captured using full observation noise covariance matrices, *R*_1_ and *R*_2_). The covariance matrix 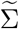 still decomposes into across- and within-area components; however, covariances at non-zero time lags (i.e., the covariance between neural activity at a time point *t*_1_ and a different time point *t*_2_ ≠ *t*_1_, indicated by the white-shaded blocks of 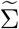 in Fig. S1b) are all zero, by definition. Just like the DLAG case (Fig. S1a), only the across-area parameters can explain across-area covariance, and within-area components fully overlap across-area components in the within-area blocks of 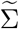 (to understand why this covariance structure is important, see case below). Across-area activity is successfully isolated by across-area variables.

The problematic case arises when we use a time series model to describe across-area interactions, but use a static model to describe within-area interactions (Fig. S1c). For example, what if we proposed a version of DLAG that simply adopted the same observation model as pCCA (i.e., full observation noise covariance matrices, *R*_1_ and *R*_2_) to model within-area interactions? In this case, although the within-area model components do explain covariance among neurons within each area, they fail to capture any within-area covariance across time points, by definition. This shortcoming forces the across-area variables to explain within-area covariance across time points. Visually, all within-area blocks of the covariance matrix 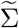 representing relationships across time points have solely magenta shading (these problematic blocks are highlighted by the ‘*’ symbols in Fig. S1c). In contrast, the true DLAG model and fully static models avoid this pitfall. These successful models (Fig. S1a,b) do not have any blocks of 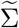 for which across-area parameters are solely responsible for explaining within-area covariance. This statistical phenomenon applies to any multi-area time series method, and is not specific to DLAG [34, 61].

---

**Figure S1.**
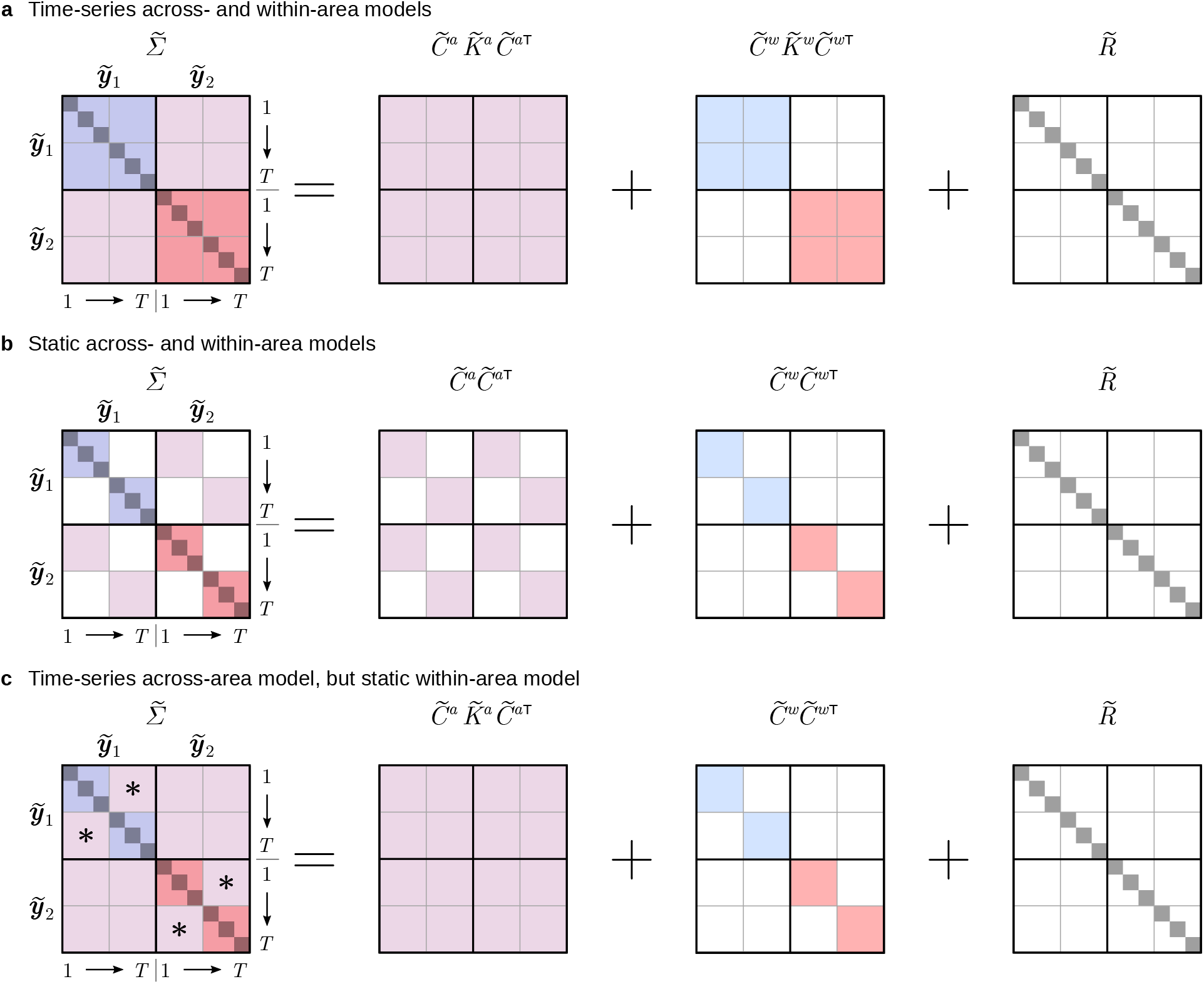
Full-sequence (trial) covariance matrix decompositions. For simplicity, in (a)-(c), we illustrate a covariance matrix for areas with three neurons each, over two time points. From left to right, panels represent the overall covariance matrix, its across-area component, its within-area component, and a component representing variance independent to each neuron. Across-area parameters (magenta shading) are solely responsible for explaining across-area covariance over time (i.e., there is no overlap of magenta with blue, red, or gray in the across-area off-diagonal blocks of the overall covariance matrix, on the left). (**a**) DLAG decomposes the covariance of a full sequence (trial) into low-rank components. Covariance among neurons within an area that cannot be explained by across-area covariance is captured by within-area parameters (area A: blue; area B: red). (**b**) Models such as probabilistic canonical correlation analysis (pCCA), for example, similarly decompose the overall covariance matrix into across- and within-area components, but make no attempt to model covariance across time points, either across or within areas (indicated by blocks with white shading). (**c**) If one is using a time series across-area model, then in the absence of a time series within-area model, across-area parameters are forced to explain within-area covariance over time. This problem is illustrated by the within-area blocks of the overall covariance matrix that have only magenta shading (indicated by the ‘*’ symbols).

